# Competition between bridged dinucleotides and activated mononucleotides determines the error frequency of nonenzymatic RNA primer extension

**DOI:** 10.1101/2021.01.02.425068

**Authors:** Daniel Duzdevich, Christopher E. Carr, Dian Ding, Stephanie J. Zhang, Travis S. Walton, Jack W. Szostak

## Abstract

Nonenzymatic copying of RNA templates with activated nucleotides is a useful model for studying the emergence of heredity at the origin of life. Previous experiments with defined-sequence templates have pointed to the poor fidelity of primer extension as a major problem. Here we examine the origin of mismatches during primer extension on random templates in the simultaneous presence of all four 2-aminoimidazole-activated nucleotides. Using a deep sequencing approach that reports on millions of individual template-product pairs, we are able to examine correct and incorrect polymerization as a function of sequence context. We have previously shown that the predominant pathway for primer extension involves reaction with imidazolium-bridged dinucleotides, which form spontaneously by the reaction of two mononucleotides with each other. We now show that the sequences of correctly paired products reveal patterns that are expected from the bridged dinucleotide mechanism, whereas those associated with mismatches are consistent with direct reaction of the primer with activated mononucleotides. Increasing the ratio of bridged dinucleotides to activated mononucleotides, either by using purified components or by using isocyanide-based activation chemistry, reduces the error frequency. Our results point to testable strategies for the accurate nonenzymatic copying of arbitrary RNA sequences.

## INTRODUCTION

The emergence of RNA-based life would have required a mechanism for RNA replication without enzymes (1-7). Replication, in turn, depends on template-directed polymerization of nucleotide building blocks—a copying step during which information stored in one piece of RNA is transferred to another. Nonenzymatic primer extension has been used extensively to study template-directed polymerization (8-12). Nonenzymatic primer extension does not work with nucleoside 5’ triphosphates because the associated activation energy is too high (13), so more reactive chemistries have been developed (9,11,14-21). The most prebiotically plausible activating group for mononucleotide-based primer extension is 2-aminoimidazole (2AI) because it can be synthesized together with 2-aminooxazole, a nucleotide precursor, and can mediate the copying of short mixed sequence templates (9,22,23). Efficient primer extension with imidazole-based nucleotide activation requires the formation of a 5’-5’phospho-imidazolium-phospho bridged dinucleotide intermediate (24,25). The bridged dinucleotide binds the template adjacent to a primer, the 3’ hydroxyl of which attacks the proximal bridging phosphate and displaces an activated nucleotide as the leaving group. Consequently, nonenzymatic RNA primer extension proceeds one base at a time even though the bridged dinucleotide pathway involves two covalently bonded nucleotides.

Mapping a prebiotic route to the emergence of heredity requires an understanding of the origin and frequency of mismatches during nonenzymatic copying. A high error rate during copying steps would quickly corrupt any beneficial sequences that may have emerged from a pool of replicating RNA molecules (26,27). Nonenzymatic copying is error-prone because it relies entirely on base-pairing, without the benefit of substrate-selective enzymes and mismatch-repair machinery. Consequently, copying G/C templates with activated G and C nucleotides occurs with higher fidelity than combinations that include A/U/T (19,28-31). Furthermore, mismatches have been found to slow down the incorporation of subsequent nucleotides, which are also more likely to be mismatched (32). This stalling effect may enable correct copying to kinetically outcompete incorrect copying (27). These and many other studies have relied on templates with defined sequences and usually one nucleotide species as a reactant so that the identities of primer extension products could be readily interpreted. In experiments with multiple activated nucleotides, or even all four, templates still had to be defined so that mismatch identities could be inferred from the known template sequences (19,31,33). A more complete understanding of how sequences are copied and how errors arise demands an assay that can characterize products on random templates with all four nucleotide reactants.

We recently developed NonEnzymatic RNA Primer Extension Sequencing (NERPE-Seq) to investigate heterogeneous reaction systems containing arbitrary template sequences and reactant nucleotides (34) (Figure 1). NERPE-Seq uses a self-priming RNA hairpin with a user-defined template. For the work presented here, the template is six random bases, and the hairpin construct is incubated with a mixture of all four 2-aminoimidazole-activated RNA nucleotides (2AIrN), isolated, processed, and deep-sequenced. The sequencing data consists of each individual template-product pair because the template and primer are physically connected in the parent hairpin. A custom analysis software package is used to filter, sort, and process the raw sequencing reads to generate information about complementary (correct) and mismatched (incorrect) products and their sequence contexts (34). These experiments tell us the yields of heterogeneous reactions, encompassing all combinations of base-dependent reactivities, competitive inhibitions, and mismatches. Here we use NERPE-Seq to identify complementary products that are preferentially synthesized, their sequence contexts, and how well primer extension samples sequence space. We also identify all mismatches generated by primer extension, and their sequence contexts. We find that complementary products harbor sequence patterns consistent with a bridged dinucleotide pathway for nucleotide incorporation, whereas sequence patterns associated with mismatches are consistent with an activated mononucleotides pathway. Finally, we test how a prebiotically plausible isocyanide-based activation chemistry (25,35), which has been shown to increase the yield of bridged dinucleotides, affects the reaction.

**Figure 1.**
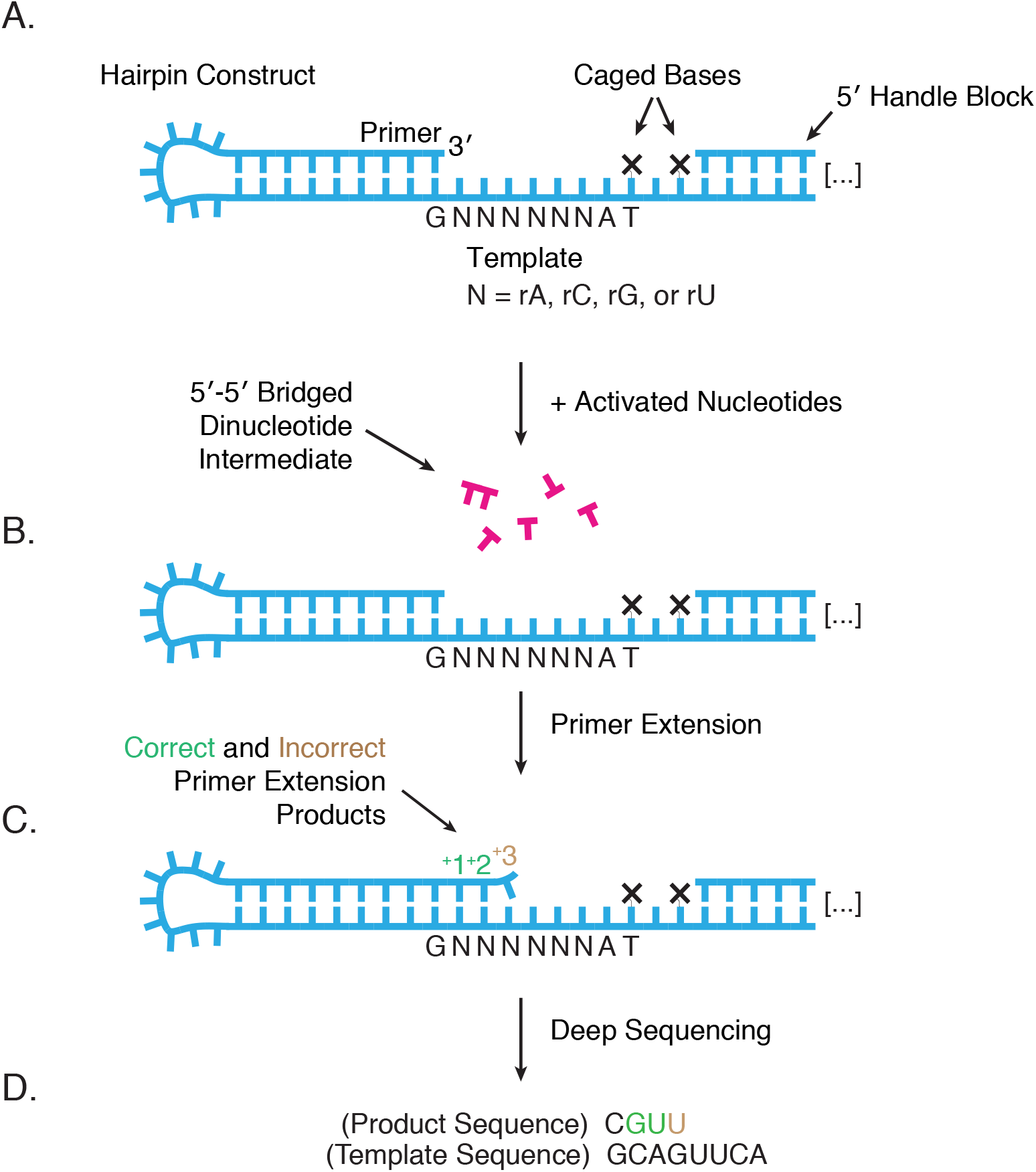
Deep Sequencing Nonenzymatic RNA Primer Extension. **A**. The RNA hairpin construct primes itself and has a six-base random-sequence template (34). The caged bases prevent primer extension from encroaching on downstream regions required for processing. Each “caged” base harbors an NPOM modification that can be removed with long-wavelength UV to generate a native dT (34,54). The 5’ Handle is required for downstream PCR, and the complementary 5’ Handle Block prevents it from interacting with the template. **B-C**. The primer is extended in the presence of all four 2-aminoimidazole-activated monoribonucleotides. The nucleotides form a reactive 5’-5’ bridged dinucleotide intermediate required for efficient copying (24). Products have both correct (complementary) and incorrect (mismatched) incorporations. **D**. Deep sequencing reports on millions of individual product-template pairs.

## MATERIAL AND METHODS

### General

Reagents, reaction conditions, syntheses, the NERPE-Seq protocol, and the data analysis code package were as previously described (34). Minor changes to the final step of sample preparation prior to sequencing submission, and the addition of a new module to the analysis code, are described below. We summarize the general procedures here in condensed form.

Bicine buffer was prepared from the Na^+^ salt and HEPES from the free acid (Sigma-Aldrich), adjusted to pH 8 with NaOH, and filtered. Enzymes and DNA/RNA ladders were purchased from New England BioLabs (NEB). Incubations at a specified temperature were performed in a Bio-Rad T100 thermal cycler. Reactions with methyl isocyanide were carried out at room temperature (∼23°C).

### Synthesis of 2-aminoimidazole-activated monoribonucleotides

0.63 mmole (1 equivalent) of the nucleoside-5’-monophosphate free acid (Santa Cruz Biotechnology or ACROS Organics) and 5 equivalents of 2-aminoimidazole hydrochloride (Combi-Blocks) were dissolved in water, the pH adjusted to 5.5 with NaOH, and the mixture lyophilized. 30 ml dry dimethyl sulfoxide (DMSO, Sigma-Aldrich) and 1.2 ml dry triethylamine (TEA, Sigma-Aldrich) were mixed under argon. To this was added the lyophilized nucleoside-5’-monophosphate and 2-aminoimidazole hydrochloride, 9 equivalents of triphenylphosphine (TPP, Sigma-Aldrich), and 10 equivalents of 2,2’-dipyridyldisulfide (DPDS, Combi-Blocks). After 30 minutes the mixture was poured into an ice-chilled solution of 400 ml acetone (Fisher Scientific), 250 ml diethyl ether (Fisher Scientific), 30 ml TEA, and 1.6 ml acetone saturated with NaClO_4_ (Sigma-Aldrich). After 30 minutes the supernatant was removed. The remaining mixture with the flocculant precipitate was centrifuged, the supernatant discarded, and the pellet washed in a solution of 1:8.3:13.3 of TEA:diethyl ether:acetone and centrifuged again. The wash was repeated twice with just acetone, and the pellets dried overnight under vacuum. Each sample was dissolved in 5 ml water, and the combined volume purified by reverse-phase flash chromatography (CombiFlash, Teledyne ISCO) over a 50 g RediSep Rf Gold C18Aq column (Teledyne ISCO) using gradient elution between (A) water and (B) acetonitrile. Target fractions were placed on ice, pooled, adjusted to pH 10 with NaOH, lyophilized, and stored at −80°C. To prepare experimentally convenient quantities, purified material was resuspended in water and the nucleotide concentration measured by a NanoDrop 2000c spectrophotometer (ThermoFisher Scientific). For experiments with all four activated nucleotides, aliquots were prepared with equimolar concentrations of each. Aliquots were lyophilized, stored at −80°C, and resuspended in water immediately prior to use.

*Oxyazabenzotriazole-activated monoribonucleotides* were prepared as in (36). Briefly, a lyophilized mixture of 0.2 g 1-hydroxy-7-aza-benzotriazole (Combi-Blocks), 0.1 g nucleoside-5’-monophosphate free acid, and TEA was resuspended in 20 ml DMSO and reacted with 1.5 g of DPDS and 1.5 g TPP. After 1-2 hours, the nucleotide was precipitated with a solution of 120 ml cold acetone, 60 ml diethyl ether, and 4.5 g sodium perchlorate. For OAtrG, the reaction mixture was left overnight, followed by a second addition of 1 g DPDS and 1 g TPP, incubated for 1-2 hours, and then precipitated. The precipitate was purified by reverse-phase flash chromatography and lyophilized.

### Synthesis and isolation of 5’-5’phospho-imidazolium-phospho bridged dinucleotide intermediates

Each bridged dinucleotide, N1*N2, was prepared by first synthesizing 2AIrN1 and OAtrN2. 2AIrN1 was then mixed with at least a 1.2x excess of OAtrN2 in water, and adjusted to pH 8.0 with NaOH/HCl. The mixture was stirred at room temperature for 1 h, then purified by reverse-phase flash chromatography using gradient elution between (A) 2 mM TEAB, pH 8, and (B) acetonitrile and lyophilized.

### Gels and gel analysis

Polyacrylamide gels (20%) were prepared with the SequaGel − Urea Gel system (National Diagnostics) and cast at 0.75 mm thick (1.5 mm for preparative) using 20 x 20 cm plates. Gels used to assay the sequencing constructs, which have no attached dyes, were removed from the glass plates and incubated in ∼140 ml of TBE + 14 μl SYBR Gold Nucleic Acid Gel Stain (Invitrogen) for several minutes then destained in TBE for ∼10 minutes. Gels were imaged with a Typhoon 9410 scanner (GE Healthcare). For analysis and visualization, TIFF-formatted images were imported into Fiji (37). Bands were quantified using the Gel Analysis function, with relative band intensities reported as the ratio of the band intensity to the total lane intensity. Band edges were excluded. Contrast and color changes were applied uniformly to the entire gel image in all cases.

### NMR measurement of bridged dinucleotide concentrations

Deuterated solvents were purchased from Cambridge Isotope Laboratories. ^1^H and ^31^P spectra were acquired on a Varian Oxford AS-400 NMR spectrometer (400 MHz for ^1^H, 161 MHz for ^31^P). Chemical shifts are reported in parts per million (ppm) on the δ scale. ^1^H NMR was referenced using the solvent resonance as the internal standard (HDO, 4.79 ppm at 25°C). All NMR spectra were recorded at 25°C. Data are reported as follows: chemical shift, multiplicity (s = singlet, d = doublet, t = triplet, q = quadruplet, m = multiplet), integration, and coupling constants. A 600 μl sample was prepared with 10 mM of each of the four 2AI-activated monomers and 200 mM deuterated Tris-Cl, pH 8. ^1^H and ^31^P spectra were acquired immediately after the sample was prepared, then at 1, 3, 6, and 24 hours. Trace TEA was used as an internal standard for integration (δ 3.20 (q, *J* = 7.3 Hz, 2H), 1.28 (t, *J* = 7.3 Hz, 3H)). ^1^H spectra were baseline corrected in MNova (Mestrelab Research), and calibrated based on TEA present in the sample from flash chromatography. The TEA concentration was determined by its integration ratio to the known concentration of mononucleotides at the initial timepoint. Activated mononucleotide and bridged dinucleotide concentrations at subsequent time points were measured by integrating corresponding ^1^H peaks. Some of the bridged dinucleotides exhibit overlapping peaks, in which case their concentrations were determined from the ^31^P spectra instead, using the peaks of other bridged dinucleotides as internal standards. Each individual bridged dinucleotide accumulates to a relatively low concentration in the 1 mM range, such that background noise contributes to the error associated with measuring their concentrations. Most of the background noise was eliminated by baseline correction. The error associated with each peak integration was calculated using the Signal-to-Noise (SNR) script in Mnova.

### Methyl isocyanide activation

The absolute concentrations of methyl isocyanide (MeNC, prepared as in (25)) and 2-methylbutyraldehyde (Tokyo Chemical Industry Co., highest available purity) were determined by comparing the integrals of their ^1^H-NMR peaks to those of adenosine-5’-monophosphate, a calibrant of known concentration. Primer extension reactions in the presence of methyl isocyanide-mediated bridge-forming activation were carried out in the same manner as for standard primer extension but with 200 mM HEPES, pH 8, as the buffer, 200 mM MeNC, 200 mM 2-methylbutyraldehyde, and 30 mM MgCl_2_.

### RNA sample preparation for deep-sequencing

Supplemental Table 1 lists oligonucleotides used in this study, and Supplemental Table 2 lists specific conditions for each deep sequencing experiment. Generally, 1 μM hairpin construct and 1.2 μM 5’ Handle Block were annealed in 200 mM buffer. Reactants and 50 mM MgCl_2_ (unless otherwise indicated) were added in a final volume of 30 μl. The mixture was incubated at 23°C, quenched, desalted, and the NPOM (caged) bases uncaged. Samples were PAGE-purified (ZR small-RNA PAGE Recovery Kit, Zymo Research). The RT Handle (template for the reverse transcription primer) was ligated with T4 RNA Ligase 2, truncated KQ. The mixture was treated with Proteinase K, phenol-chloroform extracted, and concentrated with an Oligo Clean & Concentrator spin column (Zymo Research).

The RT Primer was annealed to the purified sample and the RNA reverse transcribed with ProtoScript II. The mixture was purified with an Oligo Clean & Concentrator spin column, and the eluted cDNA stock concentration measured by spectrophotometry. 0.1 μg cDNA was added to a 50 μl Q5 Hot Start High-Fidelity DNA Polymerase PCR reaction with 0.2 μM each of NEBNext SR Primer for Illumina and NEBNext Index Primer for Illumina and run for 6 cycles with a 15 s 62°C extension step. PCR product was purified by preparative agarose gel electrophoresis (Quantum Prep Freeze ‘N Squeeze spin column, Bio-Rad) and magnetic beads (Agencourt AMPure XP, 1.7:1 volume ratio of magnetic bead suspension to sample volume). Samples were validated and concentrations measured by TapeStation (Agilent) instead of qPCR (34). Paired-end sequencing by MiSeq (Illumina, MiSeq Reagent Kit v3, 150 cycle) produced ∼20 million reads with ∼95% passing the instrument’s quality filter. The experimentally established error associated with each reported base identity is 0.062 ± 0.13% (34).

*Sequencing data analysis* was performed with the NERPE-Seq custom code package written in MATLAB (MathWorks) and described in (34). Briefly, the code filters raw data by using read quality scores, and by checking that reads agree with defined-sequence regions of the hairpin construct and that forward and reverse reads (which overlap) agree with each other. Template and product sequence pairs are extracted and characterized. The template sequence of the construct does not contain precisely equal fractions of rA, rC, rG and rU (34), so the template from a control experiment in which no activated nucleotides were added was used to generate normalization factors. All presented data are normalized— showing results as they would appear if the template were perfectly random with equal base ratios—unless otherwise indicated. A new code module reports the local sequence context of mismatches. (A key for interpreting the printed results of this module has been added to the digital resource; see link at Data Availability). The numerical data used to generate heat maps is included in the Supplementary data.

## RESULTS

### Products of primer extension copying of random sequence templates containing all four bases

#### Complementary product sequence patterns reflect reaction with the bridged dinucleotide intermediate

To begin exploring the efficiency and accuracy with which different sequences can be copied by nonenzymatic primer extension, we incubated our random-template hairpin RNA (Figure 1) with 20 mM 2AI-activated nucleotides (2AIrN) and 50 mM MgCl_2_ for 24 hours. The hairpin was then subjected to NERPE-Seq (Figure S1). Figure 2A shows the frequencies of complementary and mismatched nucleotide incorporations at each position. The reaction yield is low compared with experiments that use defined templates and reactants (9,34), but there is nonetheless a clear accumulation of products up to +3 and even +4. Increasing the concentration of 2AIrN from 20 mM to 100 mM yields more products (Figure S2A). Similarly, there are significantly fewer products in the presence of 200 mM free 2AI, which suppresses the accumulation of bridged dinucleotides (22) (Figure S2B).

**Figure 2.**
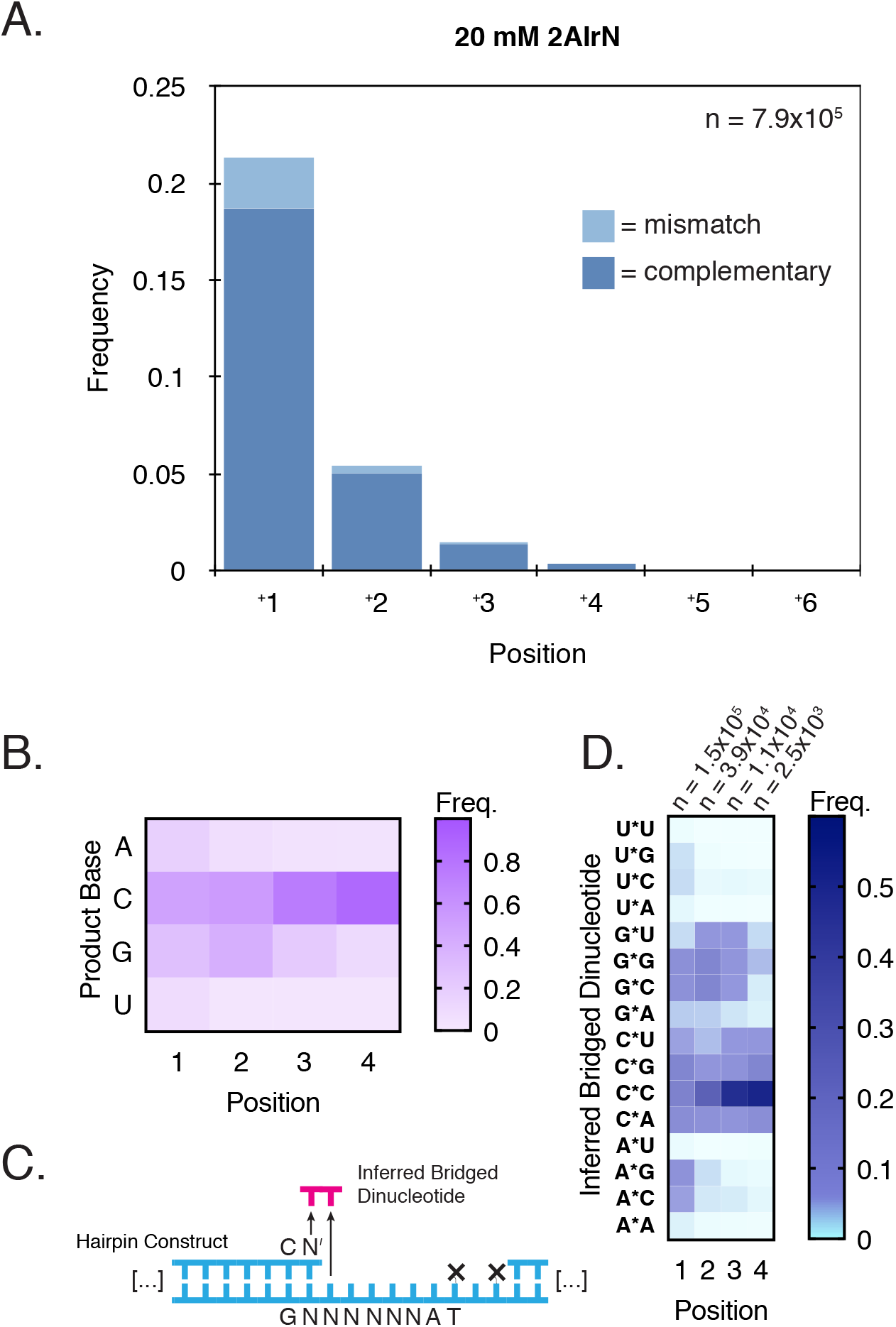
The Bridged Dinucleotide Intermediate Determines Complementary Product Sequences. **A**. Frequencies of complementary and mismatched nucleotide incorporations (20 mM 2AIrN, 24 hours; n = unextended hairpins + total nucleotide incorporation events). **B**. Position-dependent base frequencies of complementary products. **C**. Inferring bridged dinucleotide identities. The first nucleotide adjacent to the primer becomes incorporated, whereas the downstream second nucleotide functions as a leaving group an diffuses away. **D**. Position-dependent frequencies of inferred bridged dinucleotides that participated in generating complementary products.

To facilitate more detailed analyses, the data was sorted into products that are fully complementary and products with at least one mismatch. Based on prior results with defined templates, we anticipated a uniform bias in favor of primer extension with rG and rC (9). However, among fully complementary products, the incorporation of the different nucleotides was highly non-uniform. There is a marked increase in the proportion of rG and especially rC incorporations as a function of extension length (Figure 2B). The G/C-enrichment is less pronounced at position 1, which has a higher frequency of rA and rU products than downstream positions. To understand the reason for the increasing G/C bias with length, we examined the sequence context of each nucleotide incorporation event.

Because the bridged dinucleotide binds two tandem bases in the template, we examined the effect of both template positions on incorporation frequency (Figure 2C). (Recall that in the bridged dinucleotide pathway, the first nucleotide—adjacent to the extending primer—becomes incorporated as the +1 product, but the second downstream nucleotide functions as a leaving group and diffuses away.) We identified two major patterns among inferred bridged dinucleotides (Figure 2D). First, the frequencies of inferred bridged dinucleotides with a rG or rC in the first position are higher than the frequencies of those with a rA or rU in the first position, meaning that rG and rC incorporations are more common. Second, the highest frequency inferred bridged dinucleotides for each possible first position base have a rG or rC in the second position. These features are not merely due to some bridged dinucleotides forming more readily in solution than others (Figure S2C). To verify that the observed G/C bias in downstream templating positions comes from bridged dinucleotides, we performed a NERPE-Seq experiment using oxyazabenzotriazole (OAt)-based activation. OAtrNs cannot form bridged dinucleotide intermediates, and polymerize by a single-nucleotide-only pathway (38). We did not find any second-position biases in the OAt-based experiment (Figure S2D). We conclude that bridged dinucleotides with rG or rC in the second position are more likely to bind the template and react with the primer.

Consequently, products extended to some length have a high probability of harboring a downstream *templating* rC or rG. This generates a position-dependent enrichment of rC and rG template bases among extended products, explaining the patterns of terminal product bases and bases downstream of terminal products (Figure 2B, Figure S2E). The simplest explanation for these observations is that the relative affinities of bridged dinucleotides determine the templates on which primer extension proceeds.

#### Inferred bridged dinucleotide frequencies and template composition change with time and determine the composition of product sequences

We next considered how bridged dinucleotide frequencies, template composition, and complementary product sequences are related (Figure 3). We began by asking whether the inferred bridged dinucleotide frequencies could be explained by known base pairing and stacking preferences, so we turned to the empirically-derived Nearest Neighbor DataBase (NNDB) which catalogs the energetic stabilities of adjacent pairs of nucleotides (39). Our sequencing analysis reports cumulative yields and depends on out-of-equilibrium reactions, whereas the NNDB reports free energy changes associated with annealing. Nonetheless, if the various bridged dinucleotides are in comparable concentrations (Figure S2C) and competing for sites on a randomized template based on their intrinsic binding affinities, then we expect a correlation between their inferred frequencies derived from the sequencing analysis and their predicted equilibrium binding constants. We therefore plotted the frequencies of inferred bridged dinucleotides at position 1 (20 mM 2AI, 24 hour incubation) against their corresponding predicted equilibrium constants (Figure 3A). The inferred bridged dinucleotides cluster into four groups: combinations of rA and rU; combinations of rA/rU and rG/rC; A*G, C*A, G*G, and G*C; and finally C*G and C*C, with C*C having both the highest measured frequency and the highest predicted equilibrium constant. Overall, we find an R^2^ correlation coefficient of 0.76 between inferred bridged dinucleotide frequency and estimated *K*_d_. We conclude that the binding affinities of bridged dinucleotides are primarily responsible for driving complementary product formation.

**Figure 3.**
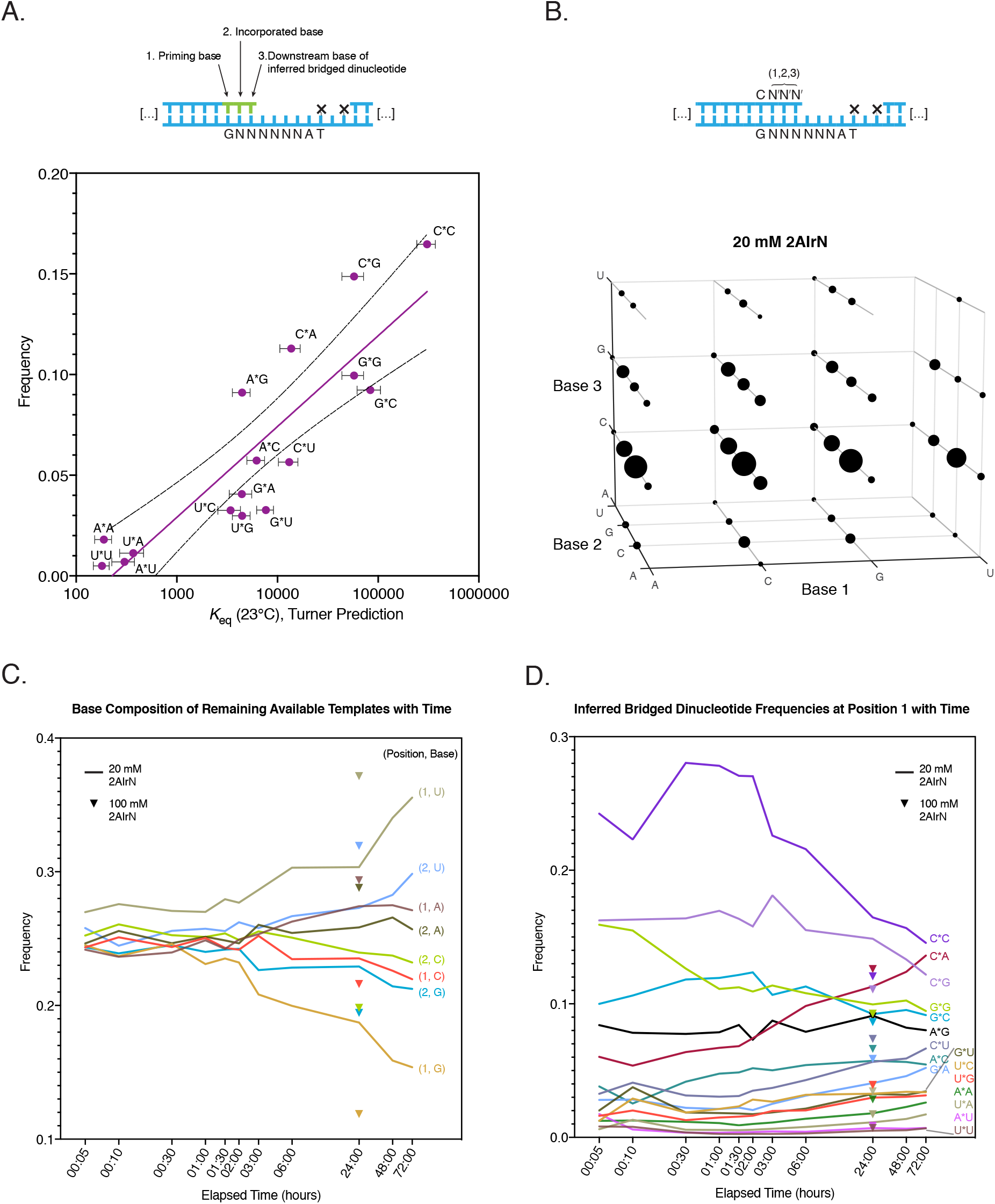
Inferred Bridged Dinucleotide Frequencies and Complementary Product Sequence Space. **A**. A log-linear correlation between predicted equilibrium binding constants and inferred bridged dinucleotide frequencies (20 mM 2AIrN, 24 hour incubation, position 1; least squares unconstrained linear fit, R^2^ = 0.76, dashed lines indicate 95% confidence interval on the fit). The Nearest Neighbor Database (NNDB, (39)) was used to calculate predicted equilibrium binding constants for each stretch of priming base, correctly incorporated base, and downstream base of a bridged dinucleotide. These values are shown plotted against the measured frequencies of inferred bridged dinucleotides at position 1 (Figure 2D). **B**. The sequence space of primer extension with 20 mM 2AIrN, 24 hour incubation. The volume of each sphere is proportional to the frequency of each complementary product at least three bases long, beginning at position 1. **C**. rG and rC templates are used up by the more reactive bridged dinucleotides, leaving behind higher proportions of rA and rU templates. **D**. More reactive inferred bridged dinucleotides participating in primer extension become less frequent with time as less reactive inferred bridged dinucleotides—dominated by combinations of rA and rU—become slightly more frequent.

We next sought to assess how the time-dependent changes in the base composition of the remaining available templates—*i*.*e*.: template positions not occupied by a product nucleotide—affect inferred bridged dinucleotide frequencies. We performed a time series of the 20 mM 2AIrN experiment, taking periodic samples up to three days, and then measured the frequencies of all remaining available template bases in positions 1 and 2 at each timepoint (Figure 3C). The frequencies of available rG and rC drop as those of rA and rU rise. (This analysis is for all available remaining templates, not just templates on which some extension has already occurred and where rG and rC are enriched as described above.) Similarly, more available sites have been consumed with 100 mM 2AIrN after 24 hours than at the same timepoint with 20 mM 2AIrN because the reaction has been pushed forward. These changes are necessarily entangled with inferred bridged dinucleotide frequencies (Figure 3D): as more of the tighter binding bridged dinucleotides lead to products on rG and rC templates, those bases become less frequent among available templates, prompting reductions in the frequencies at which rC- and rG-containing bridged dinucleotides can participate in extension. There is also a slight increase in rA- and rU-containing inferred bridged dinucleotide frequencies, with C*A increasing most dramatically, probably due to its relatively high stability (Figure 3A) and the steady rise in templating rU at position 2 (Figure 3B).

Finally, we sought to visualize how these phenomena cooperate to determine which complementary product sequences are accessed by primer extension. We measured the frequencies of products at least three bases long and plotted them by base identity at positions 1, 2, and 3 (Figure 3B and Figure S3). The volume of each sphere at each coordinate is proportional to the frequency of that triplet. Most triplets are represented, excepting a few rA/rU-rich combinations. The most prominent feature for 20 mM 2AI is the dominance of rC in the third position, which is expected given the progressive selection of rG-templates among extended products by the bridged dinucleotides (Figure 2B and 2D). The distribution becomes more uniform with 100 mM 2AIrN and several of the triplets not accessed at 20 mM 2AI appear at low frequencies (Figure S3A). Despite the challenges of copying random templates, we see examples of 51 and 62 out of 64 possible triplet sequences having been copied at 20 mM and 100 mM 2AIrN, respectively.

### Mismatches

#### Mismatch frequencies depend on position-dependent changes in template composition

Having characterized the properties of complementary products, we turned to examining mismatches. The frequency of mismatches relative to all incorporations is 11% with 20 mM 2AIrN and 13% with 100 mM 2AIrN (Figure 2A and Figure S2A). The proportion of complementary incorporations is much higher than that of mismatched incorporations at each position, demonstrating that in this system errors are not responsible for the majority of primer extension terminations. Although a mismatch is less likely to extend further than a correct incorporation (33), the majority of terminal base-pairs are correct (86% for 20 mM 2AIrN). This suggests that competition among reactants for binding sites (22,25,40,41) and the lower reactivities of some reactants may inhibit extension in the presence of all four activated nucleotides (9,31). Furthermore, the position-dependent increase in template G/C content correlates with an increased fidelity at these positions (Figure 2A). In position 1, rG is copied correctly at 95.1%, rC at 94.2%, rU at 88.4%, and rA at 54% (see below). In contrast, previous work has shown that with a defined G/C template and only 2AIrC and 2AIrG as reactants, rG is copied correctly at 99.9% and rC at 99.7% (34). This underscores the significant effect on fidelity of using all four nucleotides. It is likely that the copying of rC and rG in the template is more accurate than the copying of rA and rU because the G-C base-pair stability is greater than that of the A-U base-pair, G-U wobble-pairs, and of other mismatched pairings.

To understand mismatch patterns in greater detail, we measured the position-dependent frequency of each possible mismatch (Figure 4A). G-U wobble pairs are known to be the most stable mismatches (42), but rG templates at position 1 are largely copied correctly (Figure 3C). Consequently the A:G (template:product) pair is the most frequent mismatch at position 1 under these conditions. The G:U mismatch dominates downstream positions where there are fewer products overall (Figure 2A). The bridged dinucleotide mechanism also enriches extended products for downstream templating rG, explaining the prominent streak of G:U mismatches after position 1 (Figure 4A). The low frequency of G:A mismatches relative to A:G mismatches can also be explained by rG template depletion by correct rC incorporations at position 1. At downstream positions A:G and G:A take on comparable frequencies. U:G is also less common than G:U in position 1, but that seems to result from the identity of the priming base, which affects mismatch stabilities (42) and therefore incorporation preferences (Figure S4A).

**Figure 4.**
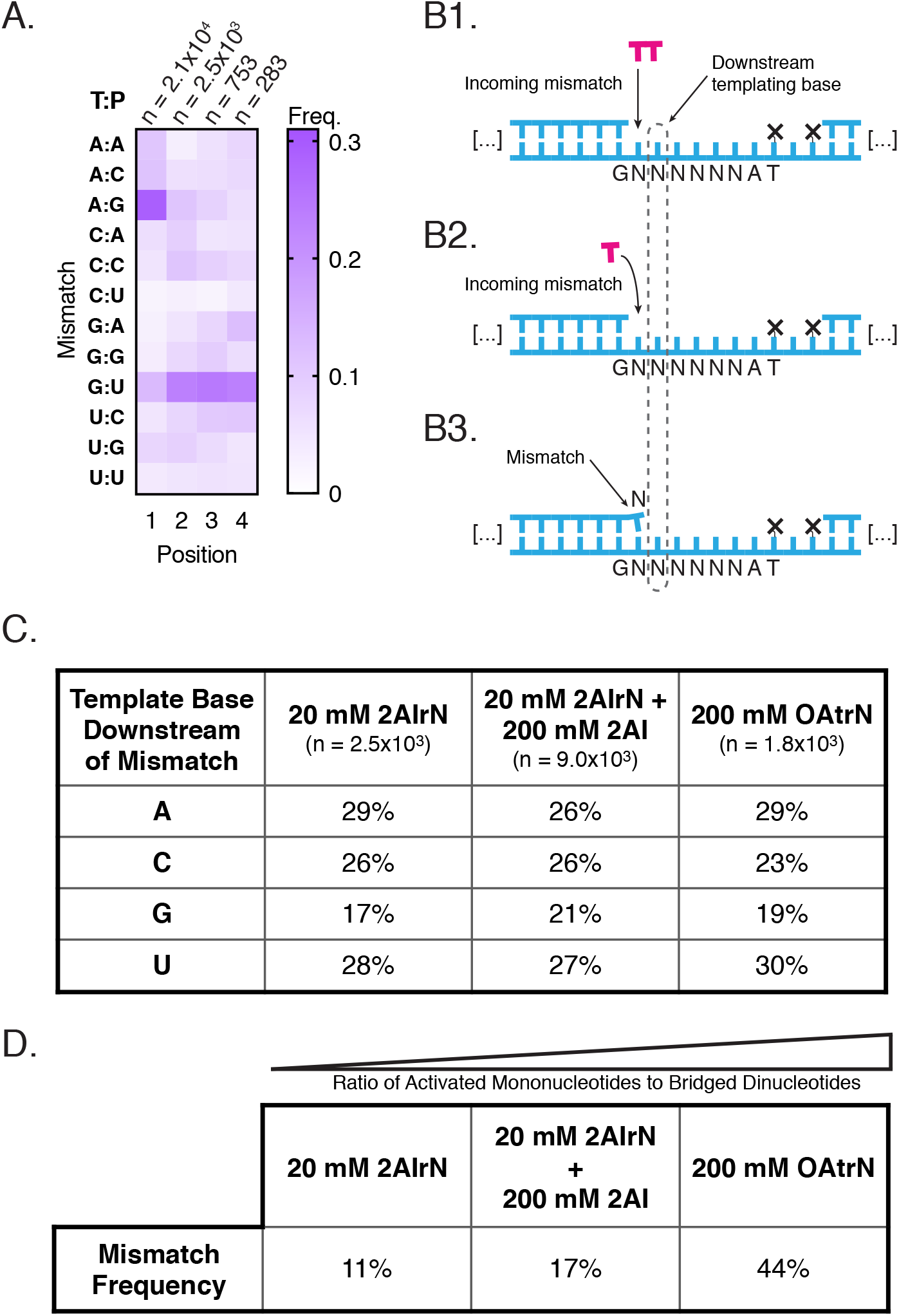
Activated Mononucleotides are Responsible for Mismatches. **A**. The position-dependent frequency of each mismatch, relative to all mismatches at the indicated position (20 mM 2AIrN, 24 hours; T:P = Template:Product). **B1-3**. Mismatches could originate from bridged dinucleotides (**B1**), or the incorporation of activated mononucleotides (**B2**). **B3**. The templating base distribution downstream of mismatches can be ascertained from the sequencing data. **C**. The templating base distribution downstream of position 1 mismatches. **D**. The overall mismatch frequency depends on the ratio of activated mononucleotides to bridged dinucleotides (increasing across experiments left to right).

NERPE-Seq also enables us to measure the effect of the incorporation of a mismatched nucleotide on subsequent incorporations. For 20 mM 2AI at 24 hours, 95% of mismatches are terminal, compared with 73% of correct incorporations. Of the mismatches that go on to prime further extension, 49% are followed by mismatches, which is much higher than the overall 11% mismatch frequency. This confirms previous observations in orthogonal systems that errors potentiate errors (27,33). Thus, the phenomenon of a cascade of errors also applies in an all-RNA, 2AI-activation based primer extension reaction with randomized components. Furthermore, mismatches that tend to be followed by correct incorporations tend to have rC as the product nucleotide, presumably because rC stacks favorably with incoming bases. In addition the two G-U wobble base-pairs, which are energetically stable and are geometrically most similar to a correct base-pair, are often followed by incorporation of a correct nucleotide (Figure S4B). Tandem mismatches show a different pattern: they are depleted in product rU and to a lesser extent rA, combinations of which do not stack as favorably as combinations of rG and rC (39,43) (Figure S4B).

#### Mismatch frequency depends on the ratio of activated mononucleotides to bridged dinucleotides

Having considered the distribution of mismatches, we next sought to discern how they form. We hypothesized that if they arise through the bridged dinucleotide pathway, then templating bases downstream of mismatches should carry a discernable signature, just as for complementary products (Figure 4B). In particular, we expected that because mismatches are less stable than correct base-pairs, the second base of any associated inferred bridged dinucleotides would be strongly enriched for rG or rC; in effect, to drive the mismatch. This would appear as a bias for templating rC and rG downstream of mismatches. However, if mismatches originate from the reaction of activated mononucleotides, there should be no such enrichment among templating bases downstream of mismatches. We therefore measured the frequencies of templating bases downstream of all mismatches (Figure 4B and 4C). Instead of a G/C bias we found roughly equal proportions of rA, rC, and rU, with rG slightly less well represented. This result suggests that mismatches arise from activated mononucleotides, and not the bridged dinucleotide.

To further test the hypothesis that mismatches arise preferentially from the reaction of a primer with an activated mononucleotide, we repeated the measurement of template bases downstream of mismatches on sequence data from the experiment with 200 mM free 2AI, which reduces the concentration of bridged dinucleotides (22). These conditions drastically reduce the amount of extended product (Figure S2B), but sufficient data was obtained to allow for an analysis of mismatch sequence context. The results show a similar unbiased distribution of template bases in the downstream position to that found above (Figure 4C). Finally, a similar result was also obtained from the experiment with 200 mM OAt-activated mononucleotides, which cannot form bridged dinucleotides. The relative frequencies of mismatches, especially at position 1, show a similar pattern to that obtained with 20 mM 2AIrN (Figure 4A and S4C, respectively). This similarity is striking given that the error frequency in the 200 mM OAtrN experiment was 44%, compared with 11% in the 20 mM 2AIrN experiment (Figure S4D and 2A, respectively). These results support our hypothesis that most mismatches result from the reaction of activated mononucleotides with the primer.

The mononucleotide model for the origin of mismatches is consistent with the observation that the mismatch frequency is correlated with the ratio of activated mononucleotide to bridged dinucleotide (Figure 4D). A salient prediction of this model is that a higher ratio of bridged dinucleotides to mononucleotides should increase fidelity. To test this, we performed experiments using purified bridged dinucleotides instead of relying on their formation *in situ*. Since bridged dinucleotides hydrolyze much more rapidly than the activated mononucleotides (22), comparisons at the 24 hour timepoint would not be useful. We therefore considered two earlier timepoints instead. After one hour the frequency of mismatches in the experiment initiated with purified bridged dinucleotides was 5.8%. Although some activated mononucleotides are expected at even early time points due to bridged dinucleotide hydrolysis, we can conclude that bridged dinucleotides contribute to mismatches at some frequency less than 5.8%. This value is still lower than the 8.5% mismatch frequency at the same timepoint under standard conditions. As expected due to further bridged dinucleotide hydrolysis, after three hours the difference shrank to 6.4% versus 7.3%. These results agree with the model that higher proportions of bridged dinucleotides lead to more accurate template copying. Finally, our time-course dataset under standard conditions, initiated with activated mononucleotides, reveals that the ratio of correct to incorrect incorporations begins low, rises for the first several hours, and then drops again (Figure S4E). A similar curve has been observed for the concentration of the bridged dinucleotide under comparable conditions (22,25). Collectively, these results show that the reaction of activated mononucleotides contributes to mismatches and that the ratio of mononucleotides to bridged dinucleotides is predictive of overall primer extension fidelity.

### Methyl isocyanide-mediated activation chemistry improves fidelity

Our model of mismatch formation prompted us to consider whether a prebiotically plausible activation chemistry could improve fidelity. Methyl isocyanide (MeNC) combined with a simple aldehyde has been found to promote nucleotide activation (35,44,45). Subsequent work with the same chemistry identified an additional pathway that promotes the formation of bridged dinucleotides from 2AI-activated mononucleotides (bridge-forming activation) (25). Bridge-forming activation exhibits several features that our model for mismatch formation predicts should reduce the frequency of errors. With bridge-forming activation, a larger fraction of the input activated mononucleotides are converted to bridged dinucleotides. Furthermore, bridge-forming activation requires less Mg^2+^, thus prolonging the lifetime of the bridged dinucleotide. We therefore incubated our random-template hairpin RNA with 10 mM 2AIrN, 30 mM MgCl_2_, and the bridge-forming activation chemistry for 24 hours. Despite the lower concentration of reactants, the product distribution is comparable to that obtained with 20 mM 2AIrN under standard conditions (Figure 5A; compare with Figure 2A). Satisfyingly, the frequency of incorrect incorporations is 7.3%, compared with 11% for the standard case. The sequence features associated with primer extension, including inferred bridged dinucleotide frequencies (Figure 5B), mismatch frequencies (Figure 5C), and complementary product sequences (Figure S5) are in agreement with those measured under standard conditions. These results further reinforce the conclusion that the ratio of activated mononucleotides to bridged dinucleotides dictates the prevalence of mismatches.

**Figure 5.**
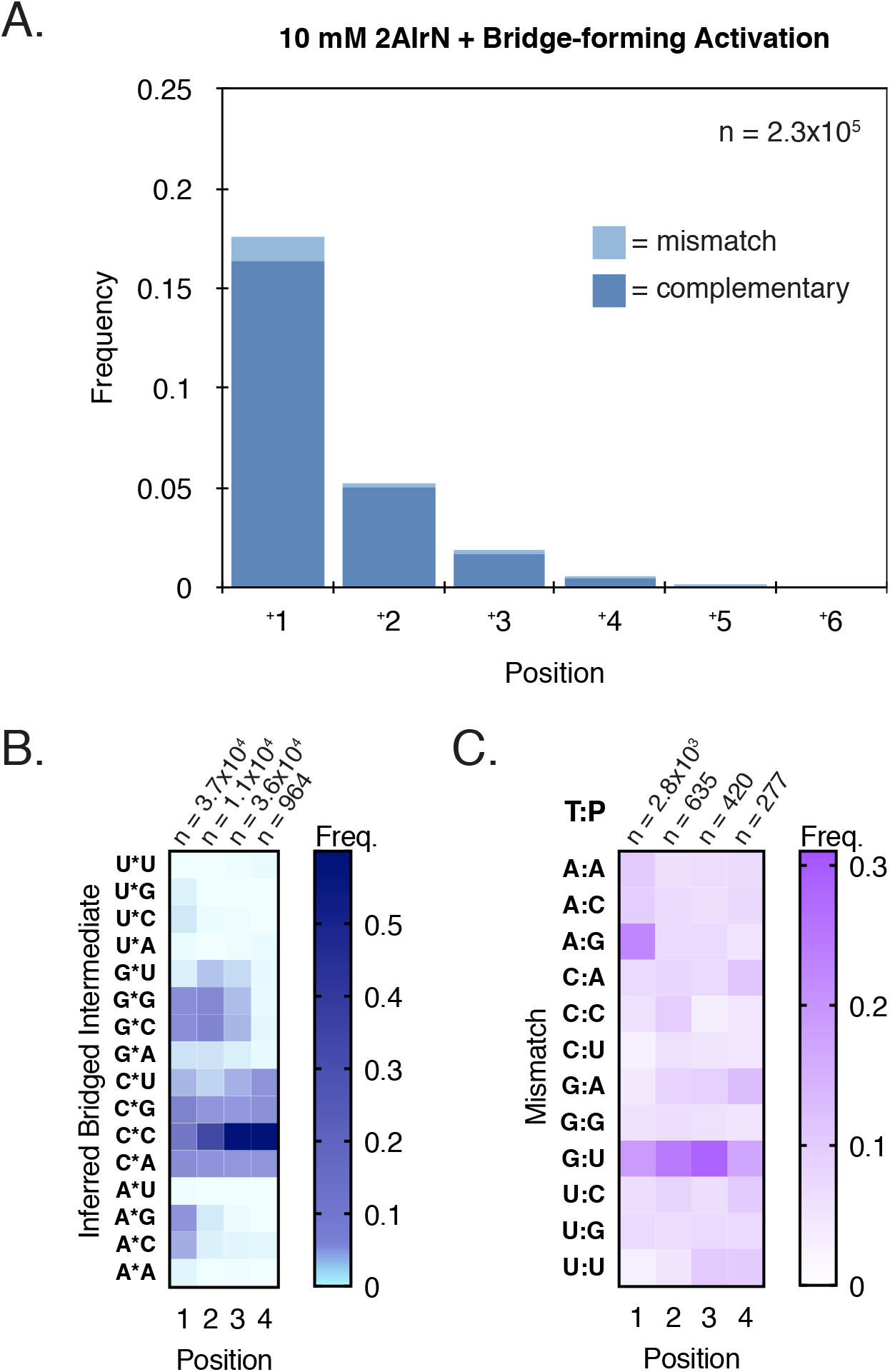
Advantages of a Prebiotically Plausible Bridge-forming Activation Chemistry. **A**. Frequencies of complementary and mismatched nucleotide incorporations (10 mM 2AIrN + MeNC-mediated bridge-forming activation, 24 hours; n = unextended hairpins + total nucleotide incorporation events). Product yields with bridge-forming activation but only 10 mM 2AIrN are comparable to that with 20 mM 2AIrN but no activation chemistry (Figure 2A), and the products are less error-prone. Other features, including bridged dinucleotide (**B**) and mismatch patterns (**C**) remain the same.

## DISCUSSION

Deep-sequencing nonenzymatic RNA primer extension enables us to measure the averaged consequences of many processes that contribute to product generation: differences in the reactivities of components, competition among them, and changes to template makeup that feed back to the behaviors of reactants. Such experiments are more challenging to analyze than those with defined templates and few reactants, but bring us a step closer to realistic scenarios, which are expected to be heterogeneous (46).

We have found that inferred bridged dinucleotide frequencies explain complementary product features, especially in selecting templates with rC or rG in the position downstream of an incorporation. This in turn affects the position- and time-dependent accumulation of products. We also discovered that the sequence pattern of downstream templating rG and rC associated with the bridged dinucleotide pathway is absent for mismatches, suggesting that mismatches arise from activated mononucleotides. An attendant insight is that some of the advantageous properties of bridged dinucleotides stem from their capacity to out-compete mononucleotides for binding sites. The two nucleotides of a bridged dinucleotide both base-pair with the template, leading to tighter binding. This contributes to the relatively low error rates observed with 2AI-based activation, even though the activated mononucleotides are at much higher concentrations than the bridged dinucleotides over the reaction time course. 2AI-activated mononucleotides are also less reactive than bridged dinucleotides (22), effectively selecting against error-prone incorporations.

Taken together, our results indicate that the ratio of bridged dinucleotides to activated mononucleotides determines the fidelity of nonenzymatic RNA primer extension, suggesting that bridged dinucleotides are more effective at directing correct incorporations. This conclusion points to several strategies for mitigating errors during nonenzymatic RNA copying of random templates. Bridged dinucleotides are more effective than activated mononucleotides at high-fidelity template copying presumably because two covalently attached nucleotides bind template sequences more selectively than individual mononucleotides. We therefore anticipate that oligos made up of two or more nucleotides should exhibit increasing selectivity for correct templates as they get longer, so that the ligation of short oligos may lead to even higher fidelity copying (10,47-50). This approach has not been attempted in a heterogeneous system, which would also contain mononucleotides to be outcompeted. Activated oligos could also serve as helpers to the mononucleotides, chaperoning them to correct templates (8). (The helper pathway is now understood to involve the formation of a bridged intermediate between a mononucleotide and a downstream oligo instead of a second downstream mononucleotide (36)).

Our results also show that fidelity with all four canonical nucleotides as templates and reactants is worse than with just rG and rC, indicating that the addition of rA and rU to the mix is responsible for the increase in overall error frequency. Previous work has identified 2-thio-uridine (s2U) as a potential alternative to rU. The s2U-rA base-pair is more stable than the rU-rA base-pair, and the s2U-rG wobble base-pair is less stable than the rU-rG wobble base-pair. These properties lead to higher fidelity copying in defined template and reactant systems where rU is replaced by s2U (31,51-53). As with ligations and helper oligos, s2U has also not been tested with random templates and a full complement of reactants. NERPE-Seq is an ideal method to examine these heterogeneous conditions.

Our sequencing assay was also found to be compatible with isocyanide-based activation chemistry, and this too promises new avenues of research. Our results show that with 2AI-based activation, some sequences are more frequent than others among complementary products, but otherwise less frequent sequences begin to appear with time or at a higher reactant concentration (Figure 3). This indicates that even in a randomized sequence context, most short sequences have the potential of appearing among products. There is no inherent reason why they cannot form. Isocyanide-based activation opens the possibility of facilitating access to these sequences by iteratively re-activating hydrolyzed bridged intermediates. Future experiments will explore the extent to which this prebiotically plausible activation chemistry combined with realistic heterogeneity will enable nonenzymatic RNA primer extension to operate with the efficiency and fidelity required for the emergence and evolution of functional sequences.

## DATA AVAILABILITY

The NERPE-Seq analysis code is available in the GitHub repository: https://github.com/CarrCE/NERPE-Seq

Raw sequencing data and NERPE-Seq analysis output data will be made available at OSF.io on formal publication.

## ACKNOWLEDGEMENTS

We thank Saurja DasGupta, Shriyaa Mittal, Aleksandar Radaković, and Marco Todisco for comments on the manuscript, and members of the Szostak laboratory for feedback and sharing experimental expertise.

## REFERENCES

1. Gilbert, W. (1986) Origin of Life - The RNA World. Nature, 319, 618–618.

2. Orgel, L.E. (1989) Was RNA the First Genetic Polymer? Evolutionary Tinkering in Gene Expression, 169, 215–224.

3. Joyce, G.F. (2002) The antiquity of RNA-based evolution. Nature, 418, 214–221.

4. Robertson, M.P. and Joyce, G.F. (2012) The Origins of the RNA World. Cold Spring Harbor Perspect. Biol., 4.

5. Krishnamurthy, R. (2015) On the Emergence of RNA. Israel Journal of Chemistry, 55, 837–850.

6. Szostak, J.W. (2017) The Narrow Road to the Deep Past: In Search of the Chemistry of the Origin of Life. Angew. Chem.-Int. Edit., 56, 11037–11043.

7. Joyce, G.F. and Szostak, J.W. (2018) Protocells and RNA Self-Replication. Cold Spring Harbor Perspect. Biol., 10.

8. Prywes, N., Blain, J.C., Del Frate, F. and Szostak, J.W. (2016) Nonenzymatic copying of RNA templates containing all four letters is catalyzed by activated oligonucleotides. eLife, 5.

9. Li, L., Prywes, N., Tam, C.P., O’Flaherty, D.K., Lelyveld, V.S., Izgu, E.C., Pal, A. and Szostak, J.W. (2017) Enhanced Nonenzymatic RNA Copying with 2-Aminoimidazole Activated Nucleotides. J. Am. Chem. Soc., 139, 1810–1813.

10. Sosson, M., Pfeffer, D. and Richert, C. (2019) Enzyme-free ligation of dimers and trimers to RNA primers. Nucleic Acids Res., 47, 3836–3845.

11. Zhou, L.J., O’Flaherty, D.K. and Szostak, J.W. (2020) Template-Directed Copying of RNA by Non-enzymatic Ligation. Angew. Chem.-Int. Edit., 59, 15682–15687.

12. Zhou, L., Ding, D. and Szostak, J.W. (2020) The Virtual Circular Genome Model for Primordial RNA Replication. Rna.

13. Rohatgi, R., Bartel, D.P. and Szostak, J.W. (1996) Kinetic and mechanistic analysis of nonenzymatic, template-directed oligoribonucleotide ligation. J. Am. Chem. Soc., 118, 3332–3339.

14. Joyce, G.F., Inoue, T. and Orgel, L.E. (1984) Non-enzymatic template-directed synthesis on RNA random copolymer: Poly(C,U) templates. Journal of Molecular Biology, 176, 279–306.

15. Joyce, G.F. and Orgel, L.E. (1986) Non-enzymatic template-directed synthesis on RNA random copolymers: Poly(C, G) templates. Journal of Molecular Biology, 188, 433–441.

16. Joyce, G.F. and Orgel, L.E. (1988) Non-enzymatic template-directed synthesis on random copolymers: Poly(C,A) templates. Journal of Molecular Biology, 202, 677–681.

17. Jauker, M., Griesser, H. and Richert, C. (2015) Copying of RNA Sequences without Pre-Activation. Angew. Chem.-Int. Edit., 54, 14559–14563.

18. Kervio, E., Sosson, M. and Richert, C. (2016) The effect of leaving groups on binding and reactivity in enzyme-free copying of DNA and RNA. Nucleic Acids Res., 44, 5504–5514.

19. Hanle, E. and Richert, C. (2018) Enzyme-Free Replication with Two or Four Bases. Angew. Chem.-Int. Edit., 57, 8911–8915.

20. Zhang, W.C., Pal, A., Ricardo, A. and Szostak, J.W. (2019) Template-Directed Nonenzymatic Primer Extension Using 2-Methylimidazole-Activated Morpholino Derivatives of Guanosine and Cytidine. J. Am. Chem. Soc., 141, 12159–12166.

21. O’Flaherty, D.K., Zhou, L.J. and Szostak, J.W. (2019) Nonenzymatic Template-Directed Synthesis of Mixed-Sequence 3 ‘-NP-DNA up to 25 Nucleotides Long Inside Model Protocells. J. Am. Chem. Soc., 141, 10481–10488.

22. Walton, T. and Szostak, J.W. (2017) A Kinetic Model of Nonenzymatic RNA Polymerization by Cytidine-5 ‘-phosphoro-2-aminoimidazolide. Biochemistry, 56, 5739–5747.

23. Fahrenbach, A.C., Giurgiu, C., Tam, C.P., Li, L., Hongo, Y., Aono, M. and Szostak, J.W. (2017) Common and Potentially Prebiotic Origin for Precursors of Nucleotide Synthesis and Activation. J. Am. Chem. Soc., 139, 8780–8783.

24. Walton, T., Zhang, W., Li, L., Tam, C.P. and Szostak, J.W. (2019) The Mechanism of Nonenzymatic Template Copying with Imidazole-Activated Nucleotides. Angew. Chem.-Int. Edit., 58, 10812–10819.

25. Zhang, S.J., Duzdevich, D. and Szostak, J.W. (2020) Potentially Prebiotic Activation Chemistry Compatible with Nonenzymatic RNA Copying. J. Am. Chem. Soc., 142, 14810–14813.

26. Eigen, M. (1971) Selforganization of matter and the evolution of biological macromolecules. Naturwissenschaften, 58, 465–523.

27. Rajamani, S., Ichida, J.K., Antal, T., Treco, D.A., Leu, K., Nowak, M.A., Szostak, J.W. and Chen, I.A. (2010) Effect of Stalling after Mismatches on the Error Catastrophe in Nonenzymatic Nucleic Acid Replication. J. Am. Chem. Soc., 132, 5880–5885.

28. Hagenbuch, P., Kervio, E., Hochgesand, A., Plutowski, U. and Richert, C. (2005) Chemical primer extension: Efficiently determining single nucleotides in DNA. Angew. Chem.-Int. Edit., 44, 6588–6592.

29. Kervio, E., Hochgesand, A., Steiner, U.E. and Richert, C. (2010) Templating efficiency of naked DNA. Proc. Natl. Acad. Sci. U. S. A., 107, 12074–12079.

30. Leu, K., Obermayer, B., Rajamani, S., Gerland, U. and Chen, I.A. (2011) The prebiotic evolutionary advantage of transferring genetic information from RNA to DNA. Nucleic Acids Res., 39, 8135–8147.

31. Heuberger, B.D., Pal, A., Del Frate, F., Topkar, V.V. and Szostak, J.W. (2015) Replacing Uridine with 2-Thiouridine Enhances the Rate and Fidelity of Nonenzymatic RNA Primer Extension. J. Am. Chem. Soc., 137, 2769–2775.

32. Rajamani, S., Ichida, J., Treco, D., Antal, T., Nowak, M., Szostak, J. and Chen, I. (2009) Non-enzymatic Primer Extension Reactions: Stalling Factors for Mismatch Extensions and Misincorporations. Origins of Life and Evolution of Biospheres, 39, 301–302.

33. Leu, K., Kervio, E., Obermayer, B., Turk-MacLeod, R.M., Yuan, C., Luevano, J.M., Chen, E., Gerland, U., Richert, C. and Chen, I.A. (2013) Cascade of Reduced Speed and Accuracy after Errors in Enzyme-Free Copying of Nucleic Acid Sequences. J. Am. Chem. Soc., 135, 354–366.

34. Duzdevich, D., Carr, C.E. and Szostak, J.W. (2020) Deep sequencing of non-enzymatic RNA primer extension. Nucleic Acids Res., 48, e70.

35. Mariani, A., Russell, D.A., Javelle, T. and Sutherland, J.D. (2018) A Light-Releasable Potentially Prebiotic Nucleotide Activating Agent. J. Am. Chem. Soc., 140, 8657–8661.

36. Walton, T., Pazienza, L. and Szostak, J.W. (2019) Template-Directed Catalysis of a Multistep Reaction Pathway for Nonenzymatic RNA Primer Extension. Biochemistry, 58, 755–762.

37. Schindelin, J., Arganda-Carreras, I., Frise, E., Kaynig, V., Longair, M., Pietzsch, T., Preibisch, S., Rueden, C., Saalfeld, S., Schmid, B. et al.. (2012) Fiji: an open-source platform for biological-image analysis. Nature Methods, 9, 676–682.

38. Sosson, M. and Richert, C. (2018) Enzyme-free genetic copying of DNA and RNA sequences. Beilstein Journal of Organic Chemistry, 14, 603–617.

39. Turner, D.H. and Mathews, D.H. (2010) NNDB: the nearest neighbor parameter database for predicting stability of nucleic acid secondary structure. Nucleic Acids Res., 38, D280–D282.

40. Deck, C., Jauker, M. and Richert, C. (2011) Efficient enzyme-free copying of all four nucleobases templated by immobilized RNA. Nat. Chem., 3, 603–608.

41. Vogel, S.R. and Richert, C. (2007) Adenosine residues in the template do not block spontaneous replication steps of RNA. Chem. Commun., 1896–1898.

42. Mathews, D.H., Sabina, J., Zuker, M. and Turner, D.H. (1999) Expanded sequence dependence of thermodynamic parameters improves prediction of RNA secondary structure. Journal of Molecular Biology, 288, 911–940.

43. Hayatshahi, H.S., Henriksen, N.M. and Cheatham, T.E. (2018) Consensus Conformations of Dinucleoside Monophosphates Described with Well-Converged Molecular Dynamics Simulations. Journal of Chemical Theory and Computation, 14, 1456–1470.

44. Patel, B.H., Percivalle, C., Ritson, D.J., Duffy, C.D. and Sutherland, J.D. (2015) Common origins of RNA, protein and lipid precursors in a cyanosulfidic protometabolism. Nat. Chem., 7, 301–307.

45. Sutherland, J.D. (2016) The Origin of Life-Out of the Blue. Angew. Chem.-Int. Edit., 55, 104–121.

46. Szostak, J.W. (2011) An optimal degree of physical and chemical heterogeneity for the origin of life? Philosophical Transactions of the Royal Society B-Biological Sciences, 366, 2894–2901.

47. Rohatgi, R., Bartel, D.P. and Szostak, J.W. (1996) Nonenzymatic, template-directed ligation of oligoribonucleotides is highly regioselective for the formation of 3’-5’ phosphodiester bonds. J. Am. Chem. Soc., 118, 3340–3344.

48. James, K.D. and Ellington, A.D. (1997) Surprising fidelity of template-directed chemical ligation of oligonucleotides. Chemistry & Biology, 4, 595–605.

49. Ninio, J. and Orgel, L.E. (1978) Heteropolynucleotides as Templates for Nonenzymatic Polymerizations. Journal of Molecular Evolution, 12, 91–99.

50. Lohrmann, R. and Orgel, L.E. (1979) Self-condensation of Activated Dinucleotides on Polynucleotide Templates With Alternating Sequences. Journal of Molecular Evolution, 14, 243–250.

51. Sheng, J., Larsen, A., Heuberger, B.D., Blain, J.C. and Szostak, J.W. (2014) Crystal Structure Studies of RNA Duplexes Containing s(2)U:A and s(2)U:U Base Pairs. J. Am. Chem. Soc., 136, 13916–13924.

52. Larsen, A.T., Fahrenbach, A.C., Sheng, J., Pian, J.L. and Szostak, J.W. (2015) Thermodynamic insights into 2-thiouridine-enhanced RNA hybridization. Nucleic Acids Res., 43, 7675–7687.

53. Prywes, N., Michaels, Y.S., Pal, A., Oh, S.S. and Szostak, J.W. (2016) Thiolated uridine substrates and templates improve the rate and fidelity of ribozyme-catalyzed RNA copying. Chem. Commun., 52, 6529–6532.

54. Lusic, H. and Deiters, A. (2006) A new photocaging group for aromatic N-heterocycles. Synthesis-Stuttgart, 2147–2150.

