## Supplementary data for "Competition between bridged dinucleotides and activated mononucleotides determines the error frequency of nonenzymatic RNA primer extension"

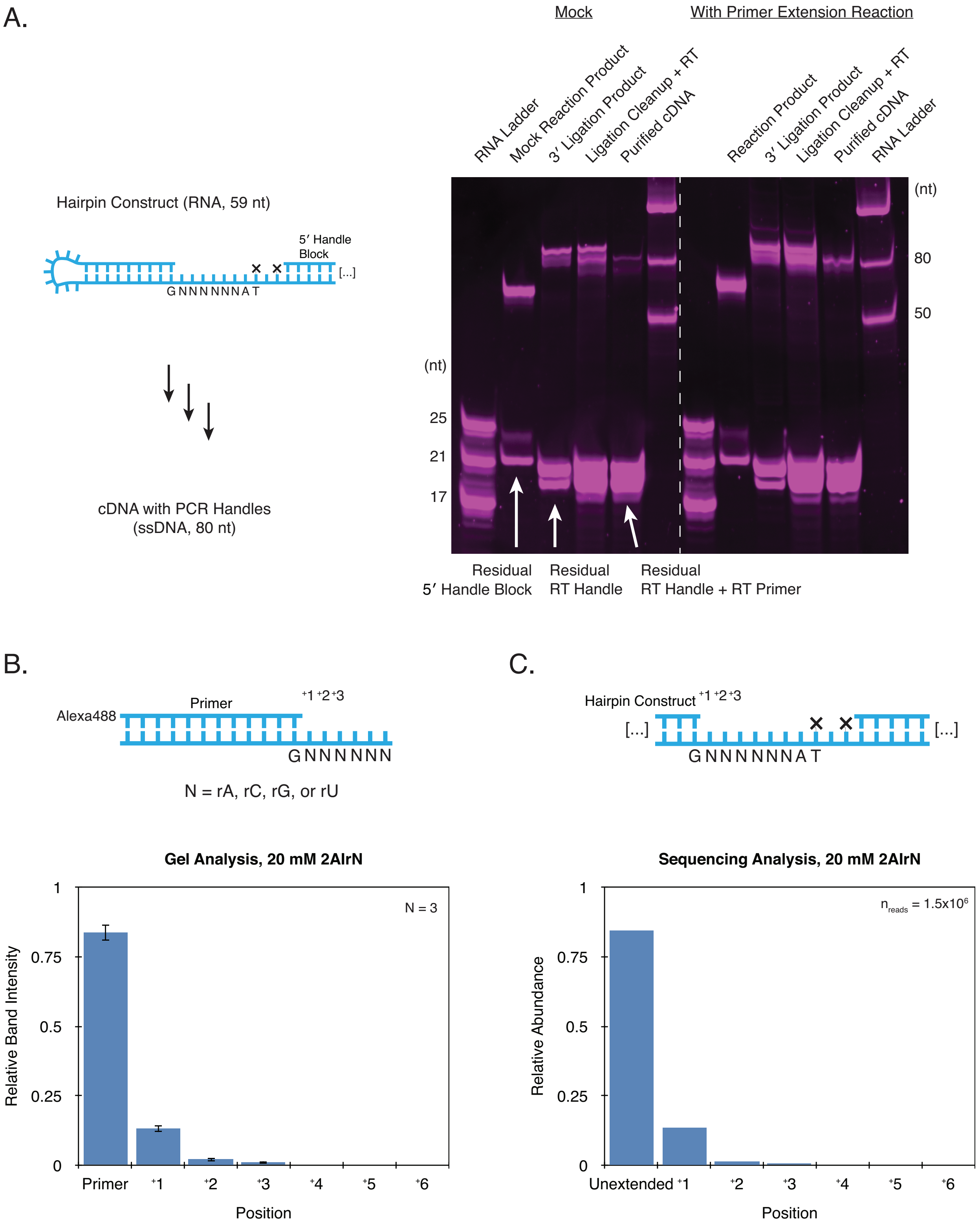
A.
Mock
With Primer Extension Reaction
RNA Ladder
Mock Reaction Product
3′ Ligation Product
Ligation Cleanup + RT
Purified cDNA
Reaction Product
3′ Ligation Product
Ligation Cleanup + RT
Purified cDNA
RNA Ladder
(nt)
80
50
(nt)
25
21
17
Hairpin Construct (RNA, 59 nt)
5′ Handle Block
[...]
GNNNNNNAT
cDNA with PCR Handles (ssDNA, 80 nt)
Residual 5′ Handle Block
Residual RT Handle
Residual RT Handle + RT Primer
B.
Alexa488
Primer
+1 +2 +3
GNNNNNN
N = rA, rC, rG, or rU
C.
Hairpin Construct
+1 +2 +3
[...]
[...]
GNNNNNNAT
Gel Analysis, 20 mM 2AIrN
N = 3
Relative Band Intensity
1
0.75
0.5
0.25
0
Primer
+1
+2
+3
+4
+5
+6
Position
Sequencing Analysis, 20 mM 2AIrN
nreads = 1.5x10^6
Relative Abundance
1
0.75
0.5
0.25
0
Unextended
+1
+2
+3
+4
+5
+6
Position

**Figure S1. Validation of the Sequencing Assay with a Random-template Hairpin Construct and all Four Activated Nucleotides. A.** Prior to deep-sequencing, the RT Handle (the template for the reverse transcription primer) is ligated to the 3′ terminus of the RNA hairpin construct, which is then reverse transcribed (RT). Denaturing polyacrylamide gel electrophoresis (PAGE) was used to visualize the efficiencies of RT Handle ligation and reverse transcription. The Mock condition ommitted activated nucleotides. Experimental conditions, in which the RNA hairpin was incubated with activated nucleotides for 24 hours, yield the same distribution of ligation and RT products as the Mock. This shows that there are no biases in the 3′ ligation of the RT Handle due to variable sequence ends. **B-C.** To validate the entire sequencing protocol, we compared the products of primer extension on a random template with all four activated nucleotides as measured by PAGE (N = number of PAGE experiments) to the equivalent reaction as measured by Nonenzymatic RNA Primer Extension Sequencing (NERPE-Seq) ($n_{reads}$ = number of sequencing read pairs). The histograms show the distribution of product lengths in each case and are in excellent agreement. This demonstrates that NERPE-Seq does not introduce biases due to sequence variability or mismatches during sample preparation or data analysis (1). Experimental conditions for **B**: 5 µM 6N Template, 1 µM Control Primer, 200 mM $Na^+$ bicine, pH 8 and water were heated to 85°C for 30 s, then cooled to 23°C at 0.2 °C/s; 20 mM 2AIrN and 50 mM $MgCl_2$ were added to initiate the reaction. All concentrations indicate final concentrations in a volume of 20 µl. 1 µl aliquots were quenched in 40 µl Urea Load Buffer (8.3 M urea [Sigma-Aldrich], 1.3x TBE buffer [from a 10x autoclaved stock], 75 µM bromophenol blue [Sigma-Aldrich, from a 7.5 mM stock in DMSO], 880 µM orange G [Sigma-Aldrich, from an 88 mM stock in DMSO], syringe-filtered), 5 µl of which was mixed with 1 µl of a 300 µM stock of Randomer Primer Extension Reverse Complement, heated to 95°C for 3 minutes then cooled to 25°C at 0.2 °C/s. 14 µl additional Urea Load Buffer was added and samples were subjected to denaturing PAGE at 5 W for 20 minutes, then 15 W for 1 hour. A control without activated nucleotides was included for comparison.

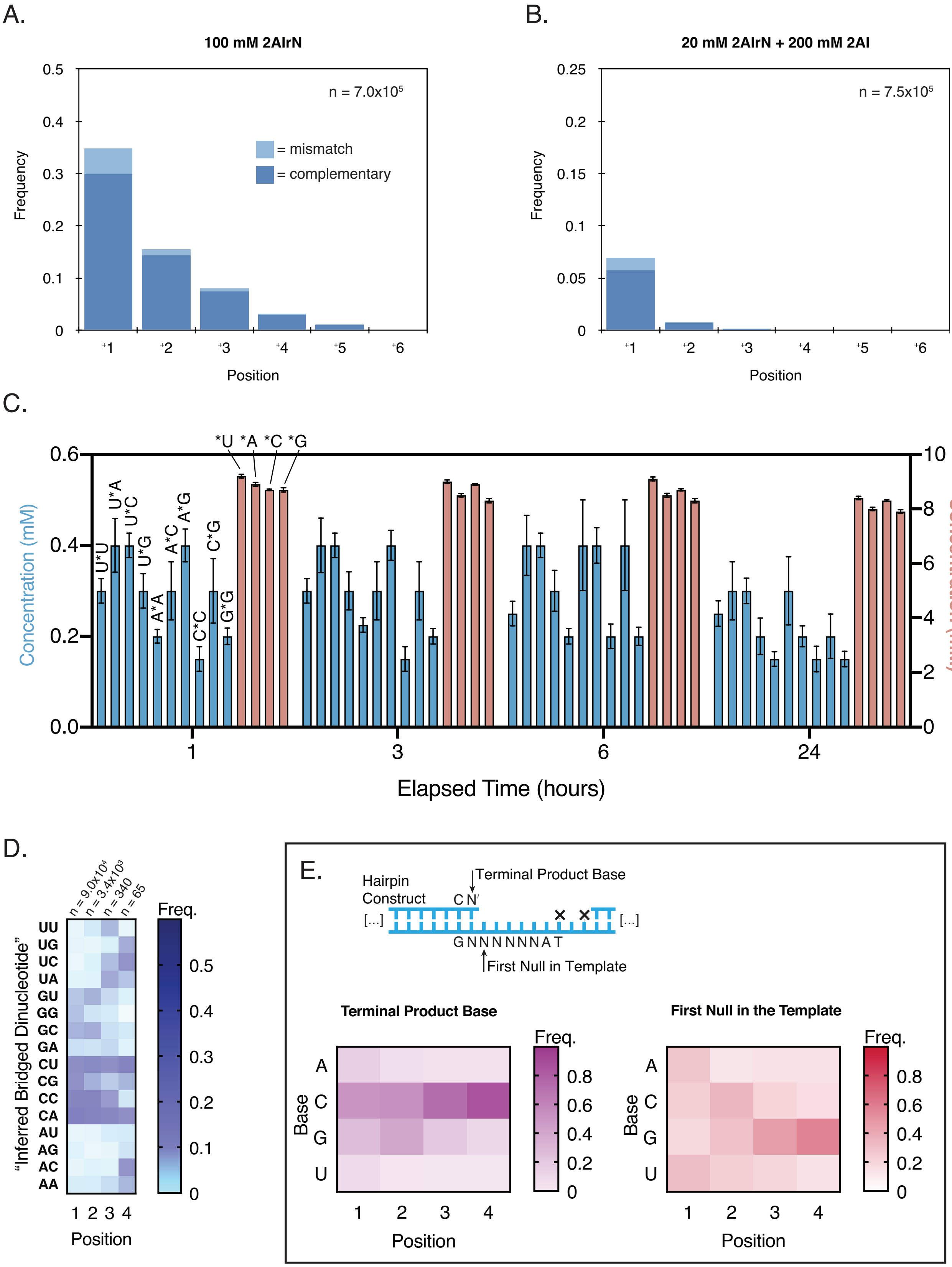
A.
100 mM 2AIrN
n = 7.0x10⁵
= mismatch
= complementary
Frequency
Position
B.
20 mM 2AIrN + 200 mM 2AI
n = 7.5x10⁵
Frequency
Position
C.
*U *A *C *G
U*U
U*A
U*C
U*G
A*A
A*C
A*G
C*C
C*G
G*G
Concentration (mM)
Concentration (mM)
Elapsed Time (hours)
D.
n = 9.0x10⁴
n = 3.4x10³
n = 340
n = 65
Freq.
"Inferred Bridged Dinucleotide"
UU
UG
UC
UA
GU
GG
GC
GA
CU
CG
CC
CA
AU
AG
AC
AA
Position
E.
Hairpin Construct
Terminal Product Base
C N′
[...]
G N N N N N N A T
First Null in Template
Terminal Product Base
First Null in the Template
Freq.
Base
Position

**Figure S2. The Bridged Dinucleotide Intermediate Determines Complementary Product Sequences, Continued. A.** Frequencies of complementary and mismatched nucleotide incorporations after 24 hours with 100 mM 2AIrN (compare with Figure 2A; n = unextended hairpins + total nucleotide incorporation events). **B.** Frequencies of complementary and mismatched nucleotide incorporations after 24 hours in the presence of 200 mM free 2AI, which inhibits accumulation of the bridged dinucleotide intermediate (compare with Figure 2A). **C.** Bridged dinucleotide (left axis scale) and activated mononucleotide (right axis scale) concentrations determined by NMR (see Material and methods for experimental details; ~10 mM starting concentration of each activated mononucleotide). Experimental error is high because the bridged dinucleotides accumulate to relatively low concentrations. Differences among many of the species are not significant, and the differences that are significant correlate with experimental error in the input concentrations of activated mononucleotides (*A and *U are at slightly higher concentrations than *C and *G). Furthermore, the differences do not correlate with inferred bridged dinucleotide frequencies. For example, C*C is measured as the lowest concentration bridged dinucleotide, but it is the most common inferred bridged dinucleotide (Figure 2D). We conclude that the concentrations of the various bridged dinucleotides are not responsible for the patterns of inferred bridged dinucleotides. ($MgCl_2$ had to be omitted from these experiments because magnesium-catalyzed hydrolysis reduces the bridged dinucleotide concentrations even further, making detection impossible.) **D.** "Inferred bridged dinucleotide analysis" of complementary products from a reaction with 200 mM OAtrN incubated for 24 hours. OAt-based primer extension cannot proceed through a bridged dinucleotide pathway; this analysis is a negative control. OAt-based primer extension is significantly less efficient than 2AI-based primer extension (see Figure S4D). Even with 200 mM reactants, very few products extend beyond +1 (note the low n-values at positions 3 and 4). **E.** Additional complementary product sequence features (data from the same experiment as in Figure 2). The distribution of terminal product bases is very similar to the overall product distribution (Figure 2B) because the majority of products of a given length are terminal (Figure S1C). The "first null in template" (see inset cartoon) distribution shows templating bases downstream of terminal product bases. There is a steady increase in rG in downstream positions because inferred bridged dinucleotides with a rC in the second position become progressively more frequent (Figure 2D).

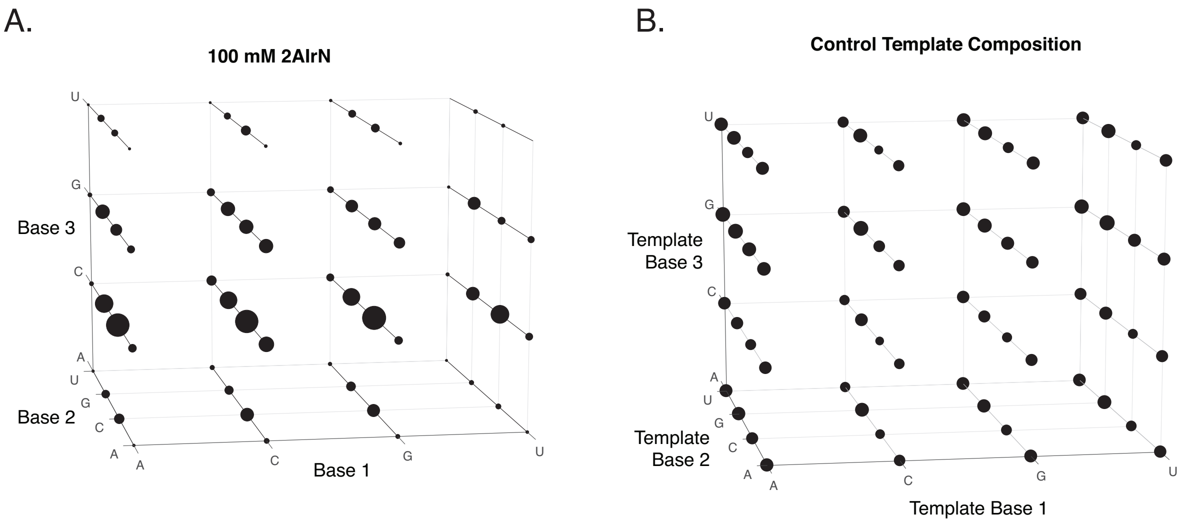


**Figure S3. Complementary Product Sequence Space. A.** The sequence space of primer extension with 100 mM 2AIrN, 24 hour incubation. The volume of each sphere is proportional to the frequency of each complementary product at least three bases long, beginning at position 1. **B.** The distribution of templating triplets in the random-template hairpin construct is relatively uniform. The data in Figure 3B and Figure S3A are not normalized because template composition biases (1) become smoothed out when considering longer stretches of templating bases.

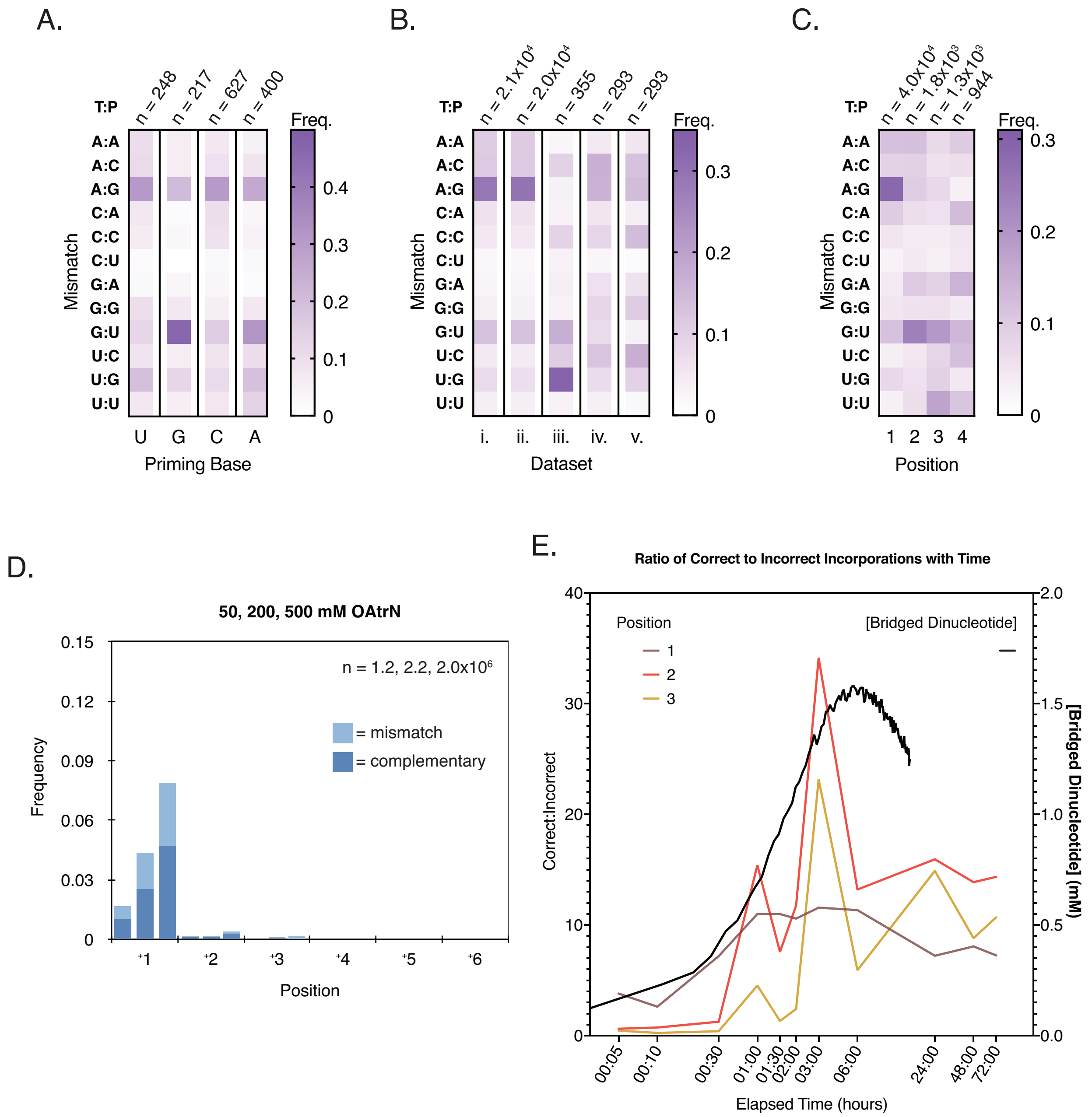


**Figure S4. Mismatch Sequence Features. A.** Mismatch frequencies at position 1 (20 mM 2AIrN, 24 hours), sorted by correctly paired priming base. This data is from an experiment with a hairpin construct that harbors all combinations of priming bases (rather than just rC, as in the primary construct [Figure 1A]). The distributions are similar except for the prevalence of G:U when the priming base is G. This combination of a GU wobble base pair adjacent to a correct base pair has been measured as the most energetically stable (2,3). (The same work also found that the G:U mismatch is always more stable than the U:G mismatch for each possible adjacent correct base pair. This explains why the frequency of G:U tends to be higher than that of U:G [Figure 4A]). **B.** Mismatch frequencies at position 1 (20 mM 2AIrN, 24 hours; T:P = Template:Product), sorted by features of adjacent positions. i. Position 1 mismatches (for comparison; same data as in Figure 4A). ii. Mismatches that are terminal. The majority of mismatches are terminal, so the distribution is the same as in (i.). iii. Mismatches that are followed by a correct incorporation. rC as the product base is favored because most of the subsequent correct incorporations are rC or rG (44% C, 38% G, 13% A, and 5.5% U), and adjacent combinations of rC and rG are energetically stable (Figure S3A). Furthermore, the inferred bridged dinucleotides that drive a correct incorporation after a mismatch skew to rC in their second positions (48% C, 19% G, 18% A, and 15% U; n = 355). G:U and U:G mismatches are probably favored because they are the most energentically similar to a correct base pair (2-4). iv.-v. Mismatches that are followed by a mismatch, and mismatches that follow a mismatch. These distributions both exhibit low frequenices of product rA and especially rU, suggesting that base stacking is important for tandem mismatch formation (5). **C.** Position-dependent mismatch frequencies with OAt-activated mononucleotides (200 mM OAtrN, 24 hours). Compare with Figure 4A. The prominant G:U "streak" found with 2AI activation is missing because without the bridged dinucleotide pathway there is no enrichment among extended products for downstream templating rG. **D.** Frequencies of complementary and mismatched nucleotide incorporations at increasing concentrations of OAtrN after 24 hours. (n = unextended hairpins + total nucleotide incorporation events.) **E.** Changes with time in the ratio of correct to incorrect incorporations (left axis) correlate with the formation of bridged dinucleotide over several hours and its subsequent hydrolysis (right axis) (6,7). The more pronounced spike in the ratio at positions 2 and 3 results from the more reactive G- and C-harboring bridged dinucleotides (Figure 2D) taking advantage of the G- and C-enriched templates selected at positions 2 and 3 by bridged dinucleotides reacting at upstream positions (Figure 3C-D). The bridged dinucleotide data is from reference (7) and shows the concentration of A*A measured by $^{31}$P NMR over 15 hours (10 mM 2AIrN, 200 mM HEPES, pH 8). The $MgCl_2$ concentration in that experiment was 30 mM instead of the 50 mM used in sequencing experiments, so the bridged dinucleotide is expected to hydrolyze more slowly. This probably explains why the bridged dinucleotide concentration peak is slightly shifted to the right relative to the correct:incorrect incorporation ratio peaks.

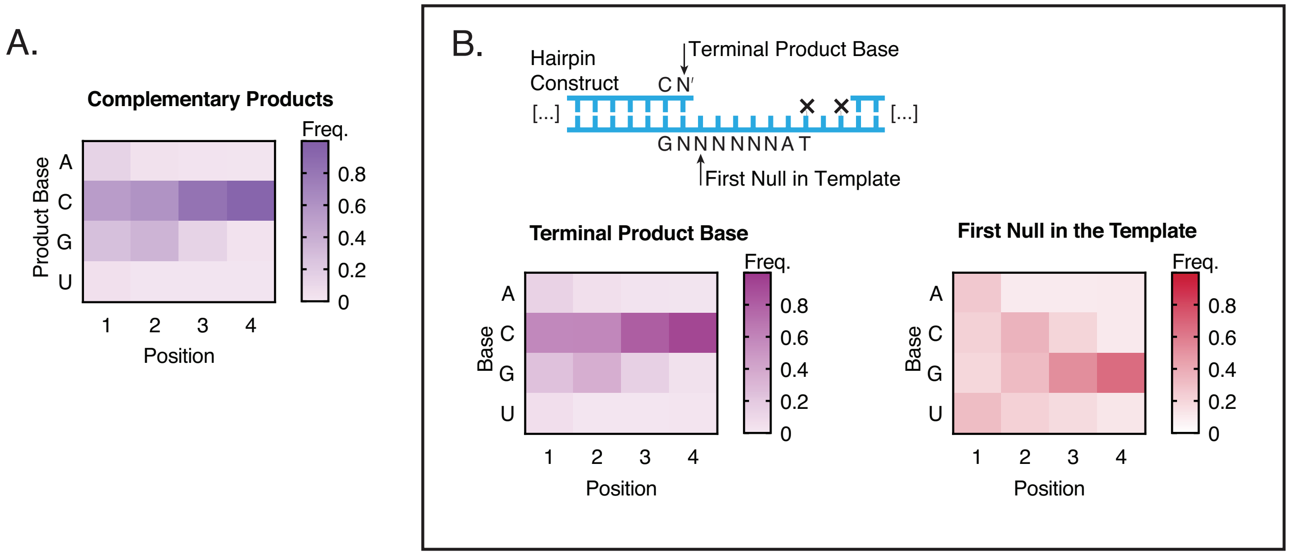


**Figure S5. Complementary Product Sequence Features with Prebiotically Plausible Bridge-forming Activation Chemistry. A.** Position-dependent base frequencies of complementary products for the same reaction as in Figure 5 (10 mM 2AIrN + MeNC-mediated bridge-forming activation, 24 hours). Compare to Figure 2B. **B.** Terminal product and first null in template distributions among complementary products for the same reaction as in Figure 5. Compare to Figure S2E.

### Supplemental Table 1, Oligonucleotides

| Oligo Name | Type | Source | Sequence (5′-3′; all termini free -OH unless otherwise noted) | Notes |
|---|---|---|---|---|
| Control Primer | RNA | IDT† | Alexa488-AGUGAGUAACUC | Figure S1 |
| 6N Template | RNA | In-house | NNNNNNGAGUUACUCACU | Figure S1 |
| Randomer Primer Extension RC‡ | RNA | In-house | AGUGAGUAACUCNNNNNN | Figure S1 |
| 6N | RNA/DNA | In-house | GUUCAGAGUUCUACAGUCCGACGAUCdT(-NPOM)CdT(-NPOM)ANNNNNNGCAUGCGACUAAACGUCGCAUGC | Random-template sequencing hairpin construct |
| MM | RNA/DNA | In-house | GUUCAGAGUUCUACAGUCCGACGAUCdT(-NPOM)CdT(-NPOM)ANNNNNNGCAUGCGACUAAACGUCGCAUGCN | Random-template sequencing hairpin construct with all possible priming base pair combinations; Figure S4A |
| 5′ Handle Block | RNA | IDT | GUCGGACUGUAGAACUCUGAA-dideoxyC | |
| RT Handle | DNA | IDT | App-AGATCGGAAGAGCACACGTCT-dideoxyC | Template for the RT Primer |
| RT Primer | DNA | IDT | AGACGTGTGCTCTTCCGATCT | |
| PCR Primer 1 (NEBNext SR Primer for Illumina) | DNA | NEB | AATGATACGGCGACCACCGAGATCTACACGTTCAGAGTTCTACAGTCCG-s-A | |
| PCR Primer 2 (NEBNext Index Primer for Illumina) | DNA | NEB | CAAGCAGAAGACGGCATACGAGAT(6-base index)GTGACTGGAGTTCAGACGTGTGCTCTTCCGATC-s-T | |

† IDT oligos ordered as RNase-free HPLC-purified
‡ RC = Reverse Complement, as competitor during PAGE analysis
⊥ See Material and Methods and (1) for details on in-house oligo synthesis and purification
App = riboA 5′-adenylation
dT(-NPOM) = NPOM-caged deoxyT (8)
N = rA, rU, rC or rG
-s- = thiol backbone linkage to inhibit exonucleases

### Supplemental Table 2, Sequencing Experiments

| Identifier | Expanded Identifier | Conditions | Notes |
|---|---|---|---|
| 6N13 | 6N hairpin construct, 1 day, #3 | 200 mM $Na^+$ bicine, pH 8, 50 mM $MgCl_2$, 20 mM 2AIrN, 24 hour incubation | Figure S1 |
| 6NC4 | 6N hairpin construct, Control, #4 | 200 mM $Na^+$ bicine, pH 8, 50 mM $MgCl_2$, 24 hour incubation | Figure S1; used as the control for 6N13 |
| 6N1 | 6N hairpin construct, 1 day | 200 mM $Na^+$ bicine, pH 8, 50 mM $MgCl_2$, 20 mM 2AIrN, 24 hour incubation | Primary experiment |
| 6NC | 6N hairpin construct, Control | 200 mM $Na^+$ bicine, pH 8, 50 mM $MgCl_2$, 24 hour incubation | Used as the control for all 6N experiments except 6N13 |
| 6NXmin | 6N hairpin construct, X = 5, 10, 30, 60, 90, 120, 180, and 360 | 200 mM $Na^+$ bicine, pH 8, 50 mM $MgCl_2$, 20 mM 2AIrN, 5-360 minute incubations | Figure 3, Figure S4E |
| 6N2 and 6N3 | 6N hairpin construct, 2 and 3 day | 200 mM $Na^+$ bicine, pH 8, 50 mM $MgCl_2$, 20 mM 2AIrN, 48 and 72 hour incubations | Figure 3, Figure S4E |
| 6N100 or 6N [100] | 6N hairpin construct, 100 mM 2AIrN | 200 mM $Na^+$ bicine, pH 8, 50 mM $MgCl_2$, 100 mM 2AIrN, 24 hour incubation | Figure 3 |
| 6NXOAt | 6N hairpin construct, X = 50, 200, and 500 mM OAtrN | 200 mM Tris-Cl, pH 9, 50 mM $MgCl_2$, 50-500 mM OAtrN, 24 hour incubation | Primer extension with OAt activation requires a higher pH than that with 2AI activation (9), Figure 4, Figure S2D, Figure S4C and D |
| 6N2AI | 6N hairpin construct, free 2AI added | 200 mM $Na^+$ bicine, pH 8, 50 mM $MgCl_2$, 20 mM 2AIrN, 200 mM 2AI (pH of stock adjusted to 8), 24 hour incubation | Figure 4, Figure S2B |
| MM20 | MM hairpin construct, 20 mM 2AIrN | 200 mM $Na^+$ bicine, pH 8, 50 mM $MgCl_2$, 20 mM 2AIrN, 24 hour incubation | Figure S4A |
| MMC | MM hairpin construct, Control | 200 mM $Na^+$ bicine, pH 8, 50 mM $MgCl_2$, 24 hour incubation | Used as the control for MM20 |
| 6NDi60 and 180 | 6N hairpin construct, Dimers, 60 and 180 minutes | 200 mM $Na^+$ bicine, pH 8, 50 mM $MgCl_2$, 2 mM each bridged dinucleotide, 60 and 180 minute incubations | |
| 6NMeNCId or 6NId | 6N hairpin construct, MeNC-based bridged activation, ideal conditions | 200 mM HEPES, pH 8, 200 mM methyl isocyanide, 200 mM 2-methylbutyraldehyde, 20 mM 2AIrN, 30 mM $MgCl_2$, 24 hour incubation | Figure 5, Figure S5 |

**Numerical Data Used to Generate each Heat-map Figure**

**Figure 2B**

| | | | | |
|---|---|---|---|---|
| A | 0.161 | 0.060 | 0.039 | 0.026 |
| C | 0.482 | 0.529 | 0.740 | 0.866 |
| G | 0.279 | 0.395 | 0.207 | 0.097 |
| U | 0.077 | 0.016 | 0.014 | 0.011 |
| | 1 | 2 | 3 | 4 |

**Figure 2D**

| | | | | |
|---|---|---|---|---|
| U*U | 0.004998 | 0.001049 | 0.000955 | 0.000556 |
| U*G | 0.029812 | 0.004021 | 0.001623 | 0.000679 |
| U*C | 0.032706 | 0.009585 | 0.010638 | 0.008816 |
| U*A | 0.011283 | 0.002273 | 0.001858 | 0.001086 |
| G*U | 0.032555 | 0.070133 | 0.061527 | 0.033267 |
| G*G | 0.099501 | 0.147142 | 0.070761 | 0.047224 |
| G*C | 0.092312 | 0.145005 | 0.064933 | 0.021964 |
| G*A | 0.040646 | 0.039765 | 0.027148 | 0.01814 |
| C*U | 0.056508 | 0.046369 | 0.061829 | 0.069822 |
| C*G | 0.148704 | 0.074235 | 0.087508 | 0.147158 |
| C*C | 0.164694 | 0.303114 | 0.492484 | 0.519812 |
| C*A | 0.112924 | 0.093359 | 0.080057 | 0.105122 |
| A*U | 0.006954 | 0.001966 | 0.00175 | 0.002921 |
| A*G | 0.091019 | 0.031064 | 0.010785 | 0.007588 |
| A*C | 0.057323 | 0.025261 | 0.022484 | 0.012432 |
| A*A | 0.018061 | 0.005659 | 0.003661 | 0.003413 |
| | 1 | 2 | 3 | 4 |

**Figure 4A**

| | | | | |
|---|---|---|---|---|
| **A:A** | **0.1077** | **0.0313** | **0.0554** | **0.0765** |
| **A:C** | **0.1130** | **0.0579** | **0.0614** | **0.0725** |
| **A:G** | **0.2909** | **0.1102** | **0.0853** | **0.0604** |
| **C:A** | **0.0608** | **0.0902** | **0.0495** | **0.0522** |
| **C:C** | **0.0512** | **0.1093** | **0.0879** | **0.0731** |
| **C:U** | **0.0215** | **0.0282** | **0.0220** | **0.0418** |
| **G:A** | **0.0272** | **0.0508** | **0.0763** | **0.1211** |
| **G:G** | **0.0344** | **0.0729** | **0.0940** | **0.0650** |
| **G:U** | **0.1298** | **0.2338** | **0.2454** | **0.2333** |
| **U:C** | **0.0470** | **0.0786** | **0.1021** | **0.1057** |
| **U:G** | **0.0764** | **0.0879** | **0.0676** | **0.0474** |
| **U:U** | **0.0400** | **0.0489** | **0.0532** | **0.0510** |
| | **1** | **2** | **3** | **4** |

**Figure 5B**

| | | | | |
|---|---|---|---|---|
| **U*U** | **0.002458** | **0.001700** | **0.004564** | **0.007918** |
| **U*G** | **0.018179** | **0.002445** | **0.000000** | **0.001936** |
| **U*C** | **0.025494** | **0.005554** | **0.004387** | **0.001477** |
| **U*A** | **0.006213** | **0.001924** | **0.003700** | **0.007733** |
| **G*U** | **0.019668** | **0.043743** | **0.033430** | **0.010531** |
| **G*G** | **0.107418** | **0.134605** | **0.047332** | **0.008409** |
| **G*C** | **0.103310** | **0.146390** | **0.049600** | **0.012747** |
| **G*A** | **0.028856** | **0.029031** | **0.020144** | **0.009397** |
| **C*U** | **0.049552** | **0.036542** | **0.051641** | **0.083575** |
| **C*G** | **0.161059** | **0.071288** | **0.060481** | **0.098127** |
| **C*C** | **0.213329** | **0.398015** | **0.600000** | **0.600000** |
| **C*A** | **0.112262** | **0.081232** | **0.068740** | **0.074589** |
| **A*U** | **0.003854** | **0.002195** | **0.001858** | **0.001665** |
| **A*G** | **0.081448** | **0.023560** | **0.005250** | **0.001965** |
| **A*C** | **0.052740** | **0.018899** | **0.014571** | **0.013282** |
| **A*A** | **0.014159** | **0.002876** | **0.003535** | **0.001389** |
| | **1** | **2** | **3** | **4** |

Figure 5C

| | | | | |
|---|---|---|---|---|
| **A:A** | **0.0960** | **0.0593** | **0.0647** | **0.0679** |
| **A:C** | **0.0921** | **0.0663** | **0.0593** | **0.0719** |
| **A:G** | **0.2224** | **0.0681** | **0.0701** | **0.0519** |
| **C:A** | **0.0691** | **0.0763** | **0.0693** | **0.1088** |
| **C:C** | **0.0556** | **0.0910** | **0.0297** | **0.0466** |
| **C:U** | **0.0269** | **0.0566** | **0.0495** | **0.0466** |
| **G:A** | **0.0402** | **0.0820** | **0.0877** | **0.1259** |
| **G:G** | **0.0538** | **0.0621** | **0.0558** | **0.0498** |
| **G:U** | **0.1862** | **0.2436** | **0.2830** | **0.1669** |
| **U:C** | **0.0606** | **0.0804** | **0.0622** | **0.0975** |
| **U:G** | **0.0693** | **0.0659** | **0.0700** | **0.0686** |
| **U:U** | **0.0278** | **0.0482** | **0.0985** | **0.0975** |
| | **1** | **2** | **3** | **4** |

Figure S2D

| | | | | |
|---|---|---|---|---|
| **UU** | **0.018066** | **0.039743** | **0.076732** | **0.017593** |
| **UG** | **0.020499** | **0.017394** | **0.031610** | **0.086030** |
| **UC** | **0.014315** | **0.029217** | **0.073746** | **0.098489** |
| **UA** | **0.023301** | **0.025531** | **0.087087** | **0.068733** |
| **GU** | **0.067223** | **0.081561** | **0.052391** | **0.029250** |
| **GG** | **0.070120** | **0.043313** | **0.036169** | **0.000000** |
| **GC** | **0.079886** | **0.085364** | **0.047649** | **0.038624** |
| **GA** | **0.047773** | **0.044577** | **0.054783** | **0.026100** |
| **CU** | **0.118651** | **0.137808** | **0.112911** | **0.156932** |
| **CG** | **0.109397** | **0.082404** | **0.063548** | **0.079286** |
| **CC** | **0.149406** | **0.151782** | **0.094752** | **0.040778** |
| **CA** | **0.183728** | **0.164051** | **0.139939** | **0.101991** |
| **AU** | **0.016627** | **0.020444** | **0.035148** | **0.036986** |
| **AG** | **0.028173** | **0.014692** | **0.020060** | **0.043671** |
| **AC** | **0.019004** | **0.027041** | **0.022965** | **0.098376** |
| **AA** | **0.033831** | **0.035077** | **0.050511** | **0.077160** |
| | **1** | **2** | **3** | **4** |

**Figure S2E, Terminal Product Base**

| | | | | |
|---|---|---|---|---|
| A | 0.153 | 0.062 | 0.043 | 0.027 |
| C | 0.508 | 0.528 | 0.725 | 0.859 |
| G | 0.256 | 0.393 | 0.216 | 0.103 |
| U | 0.082 | 0.017 | 0.015 | 0.011 |
| | 1 | 2 | 3 | 4 |

**Figure S2E, First Null in the Template**

| | | | | |
|---|---|---|---|---|
| A | 0.275 | 0.128 | 0.139 | 0.144 |
| C | 0.234 | 0.348 | 0.211 | 0.154 |
| G | 0.186 | 0.289 | 0.455 | 0.560 |
| U | 0.305 | 0.234 | 0.195 | 0.142 |
| | 1 | 2 | 3 | 4 |

**Figure S4A**

| | | | | |
|---|---|---|---|---|
| A:A | 0.1027 | 0.0592 | 0.0722 | 0.0455 |
| A:C | 0.1074 | 0.0592 | 0.0884 | 0.0812 |
| A:G | 0.3082 | 0.2039 | 0.3049 | 0.2598 |
| C:A | 0.0756 | 0.0194 | 0.1009 | 0.0335 |
| C:C | 0.0619 | 0.0290 | 0.0850 | 0.0430 |
| C:U | 0.0137 | 0.0000 | 0.0266 | 0.0239 |
| G:A | 0.0240 | 0.0338 | 0.0340 | 0.0185 |
| G:G | 0.1012 | 0.0713 | 0.0576 | 0.0723 |
| G:U | 0.1252 | 0.4652 | 0.1493 | 0.3187 |
| U:C | 0.0802 | 0.0592 | 0.0812 | 0.1036 |
| U:G | 0.1909 | 0.1237 | 0.1092 | 0.1886 |
| U:U | 0.0725 | 0.0538 | 0.0738 | 0.1328 |
| | U | G | C | A |

Figure S4B

| | | | | | |
|---|---|---|---|---|---|
| **A:A** | **0.1077** | **0.1099** | **0.0225** | **0.0494** | **0.0537** |
| **A:C** | **0.1130** | **0.1125** | **0.0951** | **0.1666** | **0.1289** |
| **A:G** | **0.2909** | **0.2977** | **0.0325** | **0.1574** | **0.1397** |
| **C:A** | **0.0608** | **0.0615** | **0.0292** | **0.0540** | **0.0404** |
| **C:C** | **0.0512** | **0.0498** | **0.0912** | **0.0900** | **0.1414** |
| **C:U** | **0.0215** | **0.0214** | **0.0292** | **0.0225** | **0.0101** |
| **G:A** | **0.0272** | **0.0266** | **0.0390** | **0.0618** | **0.0586** |
| **G:G** | **0.0344** | **0.0339** | **0.0278** | **0.0858** | **0.1071** |
| **G:U** | **0.1298** | **0.1301** | **0.1697** | **0.0755** | **0.0331** |
| **U:C** | **0.0470** | **0.0448** | **0.1043** | **0.1253** | **0.1716** |
| **U:G** | **0.0764** | **0.0717** | **0.3321** | **0.0745** | **0.0990** |
| **U:U** | **0.0400** | **0.0403** | **0.0274** | **0.0372** | **0.0165** |
| | **i.** | **ii.** | **iii.** | **iv.** | **v.** |

Figure S4C

| | | | | |
|---|---|---|---|---|
| **A:A** | **0.1178** | **0.1189** | **0.0694** | **0.0974** |
| **A:C** | **0.0850** | **0.0885** | **0.0538** | **0.0614** |
| **A:G** | **0.2832** | **0.0943** | **0.0729** | **0.0313** |
| **C:A** | **0.0958** | **0.0619** | **0.0552** | **0.1248** |
| **C:C** | **0.0473** | **0.0373** | **0.0382** | **0.0481** |
| **C:U** | **0.0259** | **0.0428** | **0.0319** | **0.0466** |
| **G:A** | **0.0400** | **0.1012** | **0.0872** | **0.1436** |
| **G:G** | **0.0481** | **0.0520** | **0.0577** | **0.0408** |
| **G:U** | **0.1160** | **0.2387** | **0.1936** | **0.1334** |
| **U:C** | **0.0362** | **0.0458** | **0.0800** | **0.1185** |
| **U:G** | **0.0720** | **0.0560** | **0.0884** | **0.0451** |
| **U:U** | **0.0326** | **0.0625** | **0.1718** | **0.1090** |
| | **1** | **2** | **3** | **4** |

Figure S5A

| | | | | |
|---|---|---|---|---|
| **A** | **0.140** | **0.043** | **0.025** | **0.019** |
| **C** | **0.531** | **0.597** | **0.826** | **0.927** |
| **G** | **0.278** | **0.349** | **0.138** | **0.038** |
| **U** | **0.051** | **0.011** | **0.012** | **0.017** |
| | **1** | **2** | **3** | **4** |

**Figure S5B, Terminal Product Base**

| | | | | |
|---|---|---|---|---|
| A | 0.1350 | 0.0488 | 0.0259 | 0.0204 |
| C | 0.5780 | 0.5820 | 0.8090 | 0.9220 |
| G | 0.2340 | 0.3570 | 0.1540 | 0.0417 |
| U | 0.0531 | 0.0117 | 0.0114 | 0.0161 |
| | 1 | 2 | 3 | 4 |

**Figure S5B, First Null in Template**

| | | | | |
|---|---|---|---|---|
| A | 0.2678 | 0.1022 | 0.1039 | 0.1090 |
| C | 0.2274 | 0.3572 | 0.2036 | 0.1084 |
| G | 0.1928 | 0.3185 | 0.5229 | 0.6595 |
| U | 0.3120 | 0.2221 | 0.1695 | 0.1232 |
| | 1 | 2 | 3 | 4 |

### SUPPLEMENTARY REFERENCES


1. Duzdevich, D., Carr, C.E. and Szostak, J.W. (2020) Deep sequencing of non-enzymatic RNA primer extension. *Nucleic Acids Res.*, **48**, e70.
2. Mathews, D.H., Sabina, J., Zuker, M. and Turner, D.H. (1999) Expanded sequence dependence of thermodynamic parameters improves prediction of RNA secondary structure. *Journal of Molecular Biology*, **288**, 911-940.
3. Turner, D.H. and Mathews, D.H. (2010) NNDB: the nearest neighbor parameter database for predicting stability of nucleic acid secondary structure. *Nucleic Acids Res.*, **38**, D280-D282.
4. Chen, J.L., Dishler, A.L., Kennedy, S.D., Yildirim, I., Liu, B., Turner, D.H. and Serra, M.J. (2012) Testing the Nearest Neighbor Model for Canonical RNA Base Pairs: Revision of GU Parameters. *Biochemistry*, **51**, 3508-3522.
5. Hayatshahi, H.S., Henriksen, N.M. and Cheatham, T.E. (2018) Consensus Conformations of Dinucleoside Monophosphates Described with Well-Converged Molecular Dynamics Simulations. *Journal of Chemical Theory and Computation*, **14**, 1456-1470.
6. Walton, T. and Szostak, J.W. (2017) A Kinetic Model of Nonenzymatic RNA Polymerization by Cytidine-5 '-phosphoro-2-aminoimidazolide. *Biochemistry*, **56**, 5739-5747.
7. Zhang, S.J., Duzdevich, D. and Szostak, J.W. (2020) Potentially Prebiotic Activation Chemistry Compatible with Nonenzymatic RNA Copying. *J. Am. Chem. Soc.*, **142**, 14810-14813.
8. Lusic, H. and Deiters, A. (2006) A new photocaging group for aromatic N-heterocycles. *Synthesis-Stuttgart*, 2147-2150.
9. Kervio, E., Sosson, M. and Richert, C. (2016) The effect of leaving groups on binding and reactivity in enzyme-free copying of DNA and RNA. *Nucleic Acids Res.*, **44**, 5504-5514.
